# CentTracker: a trainable, machine learning-based tool for large-scale analyses of *C. elegans* germline stem cell mitosis

**DOI:** 10.1101/2020.11.22.393272

**Authors:** M. Réda Zellag, Yifan Zhao, Vincent Poupart, Ramya Singh, Jean-Claude Labbé, Abigail R. Gerhold

## Abstract

Investigating the complex interactions between stem cells and their native environment requires an efficient means to image them *in situ*. *Caenorhabditis elegans* germline stem cells (GSCs) are distinctly accessible for intravital imaging; however, long-term image acquisition and analysis of dividing GSCs can be technically challenging. Here we present a systematic investigation into the technical factors impacting GSC physiology during live imaging and provide an optimized method for monitoring GSC mitosis under minimally disruptive conditions. We describe CentTracker, an automated and generalizable image analysis tool that uses machine learning to pair mitotic centrosomes and which can extract a variety of mitotic parameters rapidly from large-scale datasets. We employ CentTracker to assess a range of mitotic features in GSCs and show that subpopulations with distinct mitotic profiles are unlikely to exist within the stem cell pool. We further find evidence for spatial clustering of GSC mitoses within the germline tissue and for biases in mitotic spindle orientation relative to the germline’s distal-proximal axis, and thus the niche. The technical and analytical tools provided herein pave the way for large-scale screening studies of multiple mitotic processes in GSCs dividing *in situ*, in an intact tissue, in a living animal, under seemingly physiological conditions.

## Introduction

Tissue-resident stem cells contribute to tissue development, homeostasis and repair. To do so, stem cells rely on their ability to self-renew and differentiate into more specialized cells. Their behavior is governed *in vivo* by a specialized microenvironment, termed the niche, which provides signals that determine stem cell fate. Niche signaling protects against overgrowth or tissue loss, by controlling the size of the stem cell population and preventing inappropriate self-renewal (Januschke and Näthke, 2014; Lu and Johnston, 2013; Weaver and Cleveland, 2005). Accordingly, the interactions *in vivo* between stem cells, their niche and their tissue of residence are crucial. Live imaging approaches that permit real-time visualization of stem cells within their natural environment (i.e. intravital imaging), have provided valuable insight into stem cell biology (Park et al., 2016), yet these approaches often remain methodically challenging.

Intravital imaging of *Caenorhabditis elegans* germline stem cells (GSCs) provides an opportunity to combine well-established genetic tools with cell biological techniques to elucidate how stem cells are regulated *in vivo* (Hubbard and Schedl, 2019; Narbonne et al., 2016). *C. elegans* hermaphrodites have two U-shaped, tube-like gonad arms, each organized like a conveyer belt, with a mitotic zone at the distal end that houses the GSCs. As cells move proximally, they enter meiosis, eventually giving rise to mature gametes, which are found at the most proximal end of each gonad. GSC fate is governed by signalling from a somatic cell, termed the distal tip cell (DTC), that is located at the distal tip of each gonad arm and acts as a niche (Crittenden et al., 2006; Kimble and White, 1981). The DTC uses Notch signaling to regulate whether GSCs remain in an undifferentiated, mitotic state, or initiate differentiation by entering meiosis (Hubbard and Schedl, 2019). Like mammalian intestinal stem cells, *C. elegans* GSCs appear to self-renew according to a population model, in which GSC losses due to differentiation or damage are compensated by symmetric divisions, thus maintaining a relatively constant number of stem cells (Hubbard and Schedl, 2019; Joshi et al., 2010; Morrison and Kimble, 2006; Rosu and Cohen-Fix, 2017). Because Notch signaling relies on cell-cell interactions, maintaining contact with the niche following division could be a mechanism to control GSC fate (Kimble, 1981; Kimble and Crittenden, 2007; Kotak, 2019; Morrison and Kimble, 2006; Schofield, 1978).

Monitoring GSCs *in situ* via fluorescence microscopy is facilitated by the fact that *C. elegans* adults are transparent; however, technical challenges remain. Live imaging of dynamic subcellular events requires high temporal and spatial resolution. Notably, *C. elegans* animals are highly active and have relatively small cellular structures (e.g. the nuclei of adult GSCs are 4-5μm in diameter); thus effective, yet gentle, immobilization and higher resolution imaging methods are necessary. While several methods have been developed that allow for long-term imaging of individual *C. elegans* animals, these methods utilize a “catch and release” strategy that limits temporal resolution and is better suited for events occurring on a developmental time scale (e.g. (Guo et al., 2008; Keil et al., 2017) or rely on high-speed image acquisitions that can accommodate animal movement, but which limit spatial resolution (e.g. (Gritti et al., 2016). To image subcellular events that occur on a time scale of seconds to minutes, continuous immobilization is necessary. Continuous immobilization involves treatment with anesthetics and/or physical compression or confinement (Burnett et al., 2018; Chai et al., 2012; Hwang et al., 2014; Kim et al., 2013; Luke et al., 2014; Sulston and Horvitz, 1977), which likely limit the duration of physiologically relevant image acquisition. Recently, an elegant system was described that allows for continuous image acquisition, under seemingly physiological conditions (Berger et al., 2018); however this method relies on a fairly sophisticated microfluidic device and residual small-scale movement, such as pharyngeal pumping, may complicate automated monitoring of sub-cellular events.

We have previously described a simple mounting method that allows for high spatiotemporal imaging of mitosis in larval and adult GSCs (Gerhold et al., 2015). Here, we systematically investigated the technical factors that might impact GSC mitosis in order to define near physiological imaging conditions. We found that GSCs are sensitive to illumination intensity, anesthetic dose and physical compression, and discuss imaging parameters to minimize these effects. We further determined that preventing food intake is the major contributor to changes in GSC cell division during live imaging, and define a window of image acquisition, from the start of sample preparation to the end of visualization, during which this effect appears negligible.

To capitalize on a key advantage of working with a model system such as *C. elegans* – the ability to conduct large-scale, population-based studies – we devised a generalizable computational strategy for spatial registration, tracking and automated pairing of mitotic centrosomes, which is robust for both unperturbed and atypical GSC mitoses and which can be adapted to other cell types and model systems. We use this method to explore how mitoses and mitotic features are distributed throughout the mitotic zone and to characterize GSC mitotic spindle orientation dynamics. Our optimized live imaging conditions and largely automated image analysis thus enable large-scale studies of mitosis *in vivo*, in a highly tractable model system, facilitating the exploration of gene function within the *C. elegans* germline and, potentially, in other such complex tissues.

## Results and Discussion

### A simple mounting method for intravital imaging of GSC mitosis does not affect viability or fertility

Traditional mounting methods to immobilize *C. elegans* involve compressing the animal between an agarose pad and coverslip, with or without the addition of paralytic drugs, such as sodium azide or tetramisole (Chai et al., 2012; Fang-Yen et al., 2012; Sulston and Horvitz, 1977). The use of agarose pads micro-patterned with grooves, which approximate animal width, can constrain the worm’s typical sinusoidal body movement, thereby reducing the need for anesthetics, while also ensuring an optimal and reproducible body position (Bourdages et al., 2014; Gerhold et al., 2015; Rivera Gomez and Schvarzstein, 2018; Zhang et al., 2008). This method has the further advantage of being easy and inexpensive to implement and being suitable for use with a wide range of imaging platforms.

Our specific protocol uses a silicon wafer etched to produce several longitudinal, sloped ridges, approximately 26 μm tall and 50 μm wide, at the base, which is then used to create grooves of similar proportions in a 3% agarose pad (see Figure 2B). Worms are mounted in a minimal salt buffer (M9) supplemented with 0.04% tetramisole, gently positioned within grooves using a mouth pipet, and compressed slightly by the addition of a coverslip. While this method has been used to image *C. elegans* adults (Gerhold et al., 2015), here we focus on animals in the late L4 larval stage of development. Animals mounted in this manner, and imaged for forty minutes, showed no difference in survival or fertility when compared to controls (Figure 1A-B), indicating that this simple method has a negligible impact on viability and germline function.

**Figure 1.**
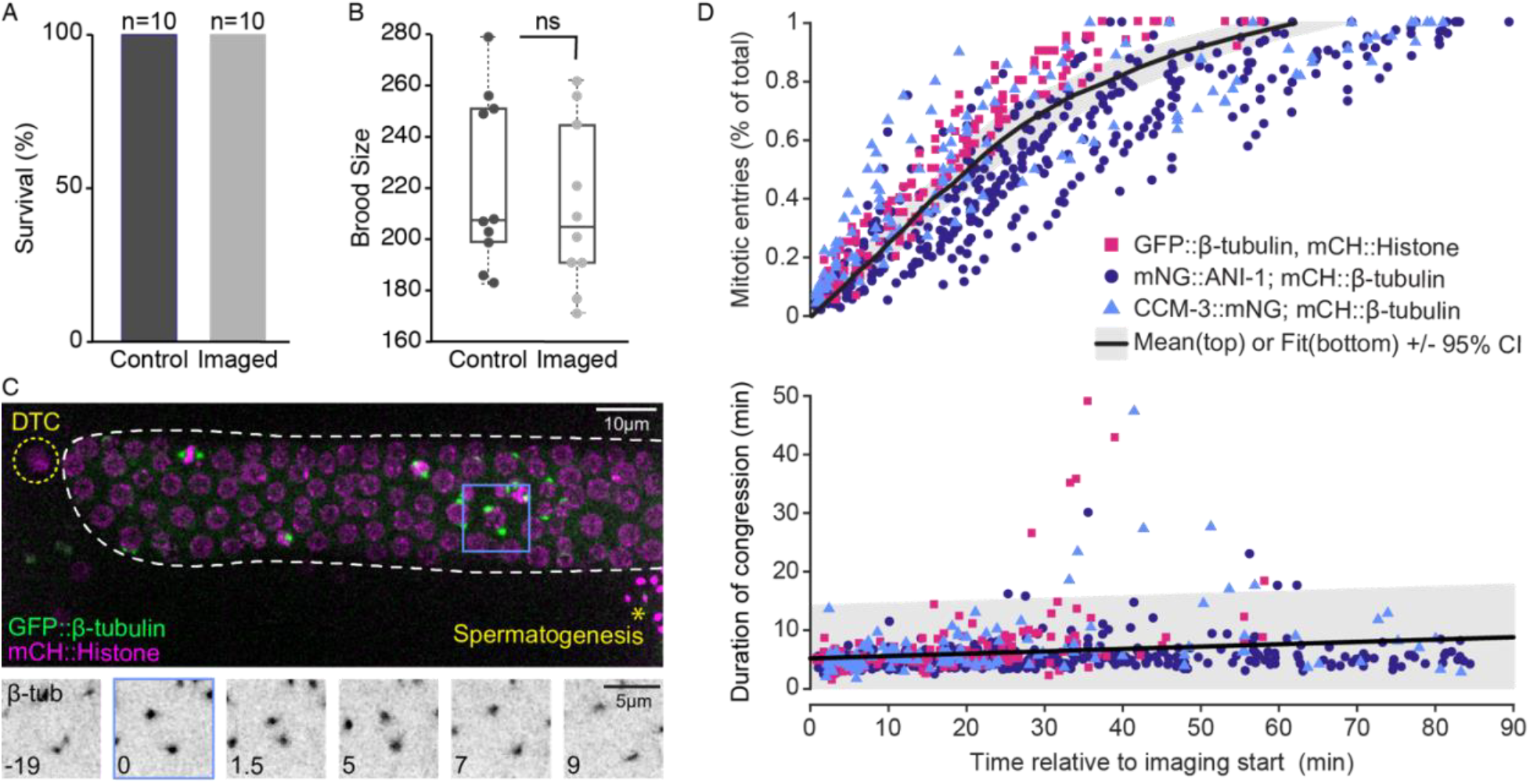
Cell cycle perturbations are observed in GSCs subjected to live imaging. (A-B) Percent survival (A) and total brood count (B) of individual animals (strain UM679: mCH::β-tubulin; mNG::ANI-1) that were left untreated (control) or imaged for 40 minutes (imaged) and recovered from the slide. Individual data points in B represent animals, box edges are the 25th and 75th percentiles, whiskers extend to the furthest data points. ns = not significant (p ≥ 0.05; two-tailed Student’s *t*-test). (C) Maximum intensity projection (top) of the distal gonad from a late L4 animal expressing GFP::β-tubulin (green) and mCH::Histone H2B (magenta) (strain = JDU19). The mitotic zone of the distal gonad is outlined in white, the nucleus of the distal tip cell (DTC) is outlined in yellow and spermatogenesis, in the proximal gonad arm, is indicated by an asterisk. Single timepoints showing a maximum intensity projection of the GFP::β-tubulin signal for the blue boxed cell are shown below. Time is in minutes relative to NEBD. (D) Mitotic entries as a percent of the total number of mitoses per individual gonad (top) and the duration of congression for each cell (bottom), over the course of 90 minute long image acquisitions, relative to image acquisition start. Data for three strains (JDU19: GFP::β-tubulin & mCH::Histone (magenta), n = 140 mitotic entries, 139 complete congressions; UM679: mCH::β-tubulin & mNG::ANI-1 (purple), n = 339 mitotic entries, 346 complete congressions; ARG16: mCH::β-tubulin & CCM-3::mNG (blue), n = 129 mitotic entries, 122 complete congressions) is shown. The mean (top) or the best fit (bottom; *m* = 0.04; r = 0.18, p = 1.203e-05) is shown in black, with the 95% confidence interval for each shaded in grey.

**Figure 2.**
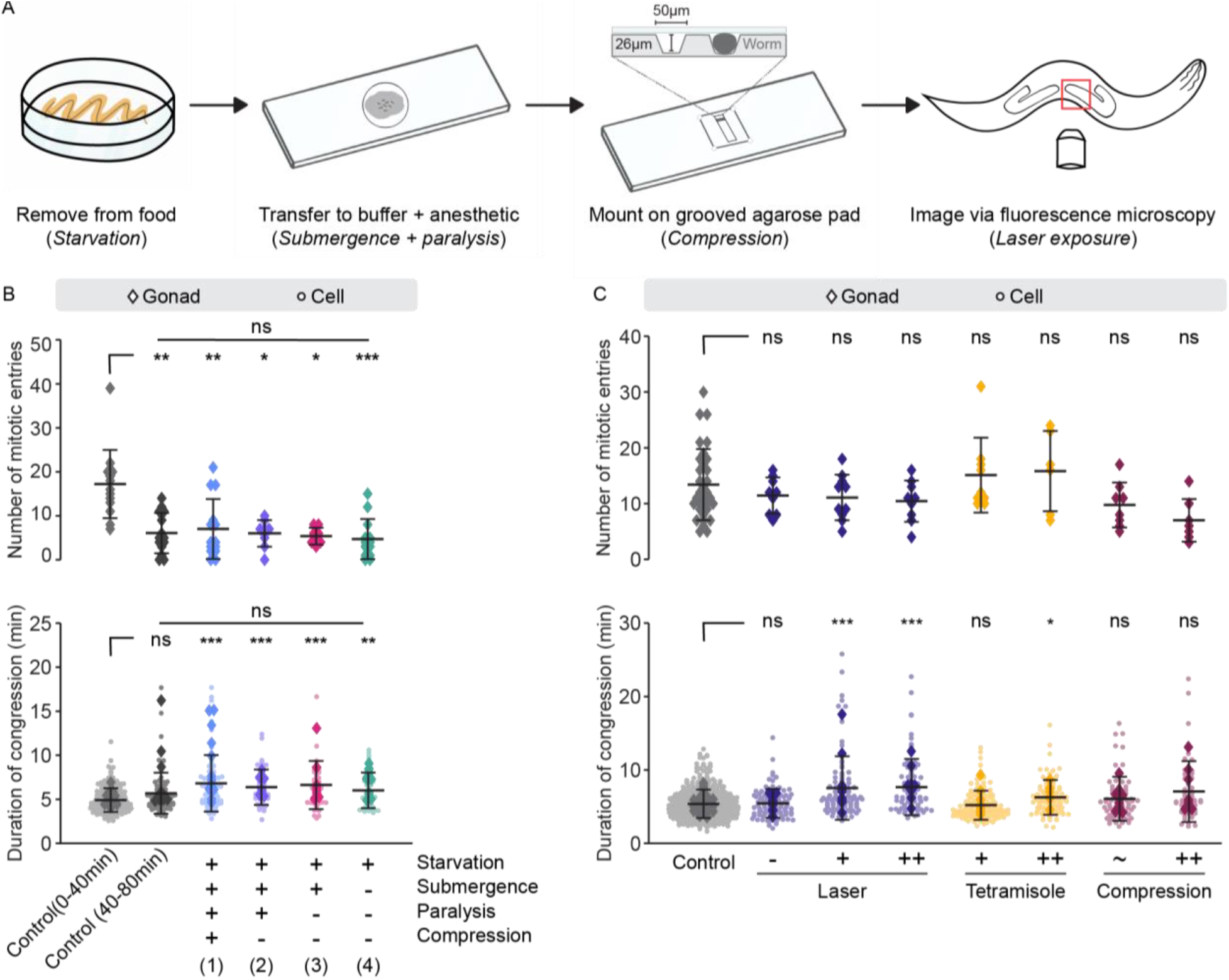
GSCs are sensitive to mounting and imaging conditions, with food deprivation being the most deleterious. (A) Schematic representation of the required steps for mounting animals and imaging GSC mitosis, with the potentially deleterious factors indicated in italics. (B) Beeswarm plots showing the number of GSCs entering mitosis per gonad (top) and the per cell (circles) and per gonad mean (diamonds) duration of congression (bottom) for mCH::β-tubulin & mNG::ANI-1 animals subjected to the indicated four treatments for 40 minutes, prior to imaging for another 40 minutes (blue, purple, magenta and green), and compared to animals imaged continuously for 90 minutes (control; grey; n = 40 animals). Control data was divided into 0-40 (n = 213 cells) and 40-80 (n = 84 cells) minute bins relative to image acquisition start. Blue = (1) fully mounted (n = 14 animals, 82 cells); Purple = (2) anesthetized in aqueous buffer (M9 with 0.04% tetramisole), but not mounted (n = 9 animals, 48 cells); Magenta = (3) submerged in aqueous buffer (M9 alone), but not anesthetized or mounted (n = 8 animals, 48 cells); and Green = (4) starved, but not submerged, anesthetized or mounted (n = 13 animals, 48 cells). (C) Beeswarm plots as in (B) for animals imaged for 40 minutes under standard conditions (grey; n = 32 animals, 396 cells), when laser intensity was modified (purple), by a 3- or 6-fold increase (+ and ++; n = 10 animals, 94 cells and n = 9 animals, 78 cells, respectively), or by eliminating the 488 laser (-; n = 9 animals, 99 cells), when tetramisole dose was increased (yellow) by 2.5-fold (+; n = 9 animals, 125 cells) or 10-fold (++; n = 6 animals, 88 cells), or when the mounting substrate was modified (magenta) by increasing the mounting pad agarose concentration to 10% with (~; n = 8 animals, 73 cells) or without (++; n = 7 animals, 48 cells) grooves. In B and C, black bars show the mean of the per animal (top) and per cell (bottom) values; error bars represent the standard deviation. Statistical analyses were done using the Kruskal-Wallis test with a Tukey-Kramer post hoc test (ns = not significant, p ≥ 0.05; * = p < 0.05; ** = p < 0.01; *** = p < 0.001). In B, p-values reflect the comparison to Control (0-40min). All other comparisons were not significant. In C, p-values reflect the comparison to Control.

### Cell cycle perturbations are observed in GSCs subjected to live imaging

To monitor GSC mitosis, we use *C. elegans* strains that express a fluorescently tagged version of β-tubulin under the control of germline-specific regulatory sequences, combined with a second marker to follow other structures of interest (e.g. nuclei; Figure 1C). We then track centrosome pairs, measure the distance between them in three dimensions (hereafter “spindle length”), and use reproducible changes in spindle length to define various mitotic landmarks, including nuclear envelope breakdown (NEBD) and anaphase onset (Figure 1C, see also Figure 4E; (Gerhold et al., 2015). To diagnose the state of GSCs during live imaging, we took advantage of the fact that cells must generally satisfy cell cycle checkpoints to undergo proper division. We assessed two checkpoints: the G2/M checkpoint, which prevents mitotic entry in response to DNA damage, stress signaling and dietary conditions (Ables et al., 2012; Rieder, 2011), and the spindle assembly checkpoint (SAC), which delays anaphase onset and mitotic exit in the presence of unattached kinetochores, the multi-protein structure responsible for connecting chromosomes to spindle microtubules (London and Biggins, 2014).

We considered NEBD as the definitive sign of mitotic entry and satisfaction of the G2/M checkpoint, and anaphase onset as an indication that cells had satisfied the SAC. To determine how long cells took to satisfy the SAC, we measured the duration of congression, which we define as the period of time after NEBD, once spindle length reaches a constant minimum, until the start of anaphase pole separation (see Figure 4E; (Gerhold et al., 2015). Over the course of ninety minute-long image acquisitions, we found that the number of cells entering mitosis decreased substantially, such that more than 80% of mitotic entries occurred within the first forty minutes of imaging and almost none were observed after sixty minutes (Figure 1D). Further, the duration of congression tended to increase slightly over the ninety-minute imaging period (Figure 1D). We conclude that GSCs are sensitive to our imaging/mounting regime, with the overwhelming effect being a decrease in the number of mitotic entries, suggesting that the cell cycle arrests at or prior to the G2/M checkpoint. Importantly, these results suggest that, under commonly used animal mounting conditions, there is a limited time window for physiologically relevant imaging of GSC mitosis.

### Food deprivation is the primary factor impacting GSCs during live imaging

During mounting and live imaging, *C. elegans* animals are exposed to several potentially deleterious factors. These include laser exposure, treatment with paralytic drugs, physical compression, submergence in an aqueous buffer and removal from a source of food and/or prevention of food intake due to paralysis (i.e. starvation; Figure 2A). As these factors are common to many mounting/imaging methods, we sought to explore how each contributes to changes in GSC cell cycle progression. To do so, we looked at the number of mitotic entries and the average duration of congression in animals that were continuously mounted and imaged for 90 minutes, and divided cells into two bins: those dividing between 0 and 40 minutes of imaging and those dividing between 40 and 80 minutes of imaging. We then compared cells in the second bin to cells from animals subjected to the following four treatments for 40 minutes, prior to being imaged for a further 40 minutes: (1) fully mounted, but not imaged; (2) anesthetized in aqueous buffer (M9 with 0.04% tetramisole), but not mounted or imaged; (3) submerged in aqueous buffer (M9 alone), but not anesthetized, mounted or imaged; and (4) transferred to a normal feeding plate without a food source, i.e. starved, but not submerged, anesthetized, mounted or imaged (Figure 2B). Following all four treatments, we found that the number of mitotic entries was decreased and the duration of congression was increased, relative to cells in the first bin (0-40 minutes of imaging) and to similar extent to what we observed in cells in the second bin (40-80 minutes of imaging). Thus removing animals from food is sufficient to produce the changes in GSC mitosis that we observe under continuous imaging conditions, suggesting that starvation is the principal perturbing factor for GSC cell cycle progression under our mounting/imaging regime.

### GSCs are sensitive to laser exposure, anesthetic dose, mounting substrate and temperature

To further investigate the impact of our mounting/imaging conditions on GSCs, we varied each factor individually to determine whether they could effect changes in GSC cell cycle progression. Effective spatial resolution is linked to the signal-to-noise ratio (SNR) of an imaged specimen (Stelzer, 1998), and while increasing sample illumination may improve SNR, this often leads to phototoxicity in living samples (Dixit and Cyr, 2003). Thus the photosensitivity of GSCs must be considered when designing and interpreting live imaging experiments. Similarly, the impact of paralytic agents used to immobilize worms and thus facilitate high resolution imaging must also be assessed. Tetramisole is a cholinergic inhibitor commonly used for immobilizing *C. elegans* (Aceves et al., 1970; Lewis et al., 1987; Thienpont et al., 1966), but its effect on GSCs has not been determined.

To test how light intensity affects GSCs, we increased the intensity of both excitation lasers by 3- or 6-fold, relative to our standard conditions, which use the minimum laser power necessary to visualize centrosomes in one channel and a second structure of interest in the other. To reduce light exposure, we imaged GSCs using only a single laser (561 nm), visualizing centrosomes alone. While neither increasing nor reducing laser exposure affected the number of mitotic entries, the duration of congression was significantly longer and more variable when laser intensity was increased (Figure 2C). To test whether the dose of tetramisole impacts dividing GSCs, we increased the concentration by 2.5- and 10-fold (0.1 and 0.4%, respectively) relative to the minimal concentration required to effectively immobilize worms for imaging (0.04%). We found that while increasing the concentration of tetramisole did not significantly affect the number of cells entering mitosis, a 10-fold increase lead to significantly longer durations of congression (Figure 2C). Thus, both laser exposure and tetramisole dose can affect GSC mitosis, largely by delaying mitotic exit; however, visualization of GSCs is possible at laser intensity settings and tetramisole doses well below the threshold at which deleterious mitotic effects are observed.

Many methods for mounting *C. elegans* animals use a relatively rigid agarose pad, which can immobilize the worm via compression after a coverslip has been applied (Dong et al., 2018). The use of micropatterned grooves that approximate animal size likely decreases this compression. Our mounting method makes use of an intermediate agarose concentration (3%), which preserves the integrity of the grooves without being overly rigid. To test whether agarose rigidity and/or compression impacts GSCs, we imaged animals on 10% agarose pads, with and without micropatterned grooves. In worms mounted on 10% agarose pads without grooves, we observed a trend towards fewer mitotic entries and longer durations of congression (Figure 2C). While neither trend was significant compared to control (p=0.0825 and p=0.094, respectively), the duration of congression was more variable under these conditions, and the frequency of overly delayed cells was higher, with 21% of cells displaying a duration of congression greater than the 95^th^ percentile value for controls. These results suggest that physical compression of the animal may perturb GSC divisions and is not an optimal mounting strategy. Notably, the mitotic parameters of GSCs imaged in animals mounted on 10% agarose pads with grooves were similar to those in animals mounted using our standard method (Figure 2C). This suggests that agarose micropatterning reduces the negative impact of physical compression and allows for more physiologically accurate observations of GSC divisions.

While grooves were typically made using custom-microfabricated silicon wafers, we also tested the effect of agarose micropatterning using vinyl Long Play (LP) records, as described previously (Rivera Gomez and Schvarzstein, 2018; Zhang et al., 2008). We found that mounting animals on 3% agarose pads molded with grooves from a vinyl record yielded reasonably good results, with a similar number of mitotic entries and a slight, but significant, increase in the average duration of congression in GSCs when compared to our silicon wafers (Figure S1A). Thus LP-molded grooves can serve as an inexpensive and readily available alternative to micro-patterned silicon wafers for GSC live imaging.

Finally, *C. elegans* is generally maintained in the lab within a temperature range of 15-25°C, and most imaging experiments are conducted at “room temperature”. However, room temperature can vary widely and may not be an accurate reflection of the actual temperature experienced by the sample during imaging. To assess the impact of temperature on GSC mitosis, we imaged animals at 15, 20 and 25°C, using a microfluidic temperature control device (CherryTemp) to maintain our samples at a stable temperature. While the number of mitotic entries appeared to be relatively constant across temperatures, GSCs from animals imaged at 15°C had significantly longer durations of congression (Figure S1B), suggesting that temperature is an additional factor that ought to be considered when live-imaging GSCs.

In sum, we find that mitotic entry and exit in GSCs are sensitive to an array of factors that are commonly used in mounting and imaging *C. elegans* animals. However, our results indicate that minimally perturbing conditions can be found to afford high spatiotemporal live imaging suitable for mitotic studies. Our results indicate that food removal is the main factor to consider when live imaging GSCs, providing further evidence that cell cycle progression in GSCs is highly responsive to dietary conditions (Gerhold et al., 2015; Hubbard et al., 2013; Seidel and Kimble, 2015). Importantly however, our analysis defines a window of roughly 40 minutes, post-food removal, during which GSC physiology, at least with respect to mitosis, is minimally perturbed.

### CentTracker enables automated monitoring and analysis of GSC mitosis

To follow GSC mitosis, we track individual centrosomes in three dimensions and manually pair centrosomes that belong to the same cell. Tracking and pairing is performed using the open-source platform TrackMate in Fiji (Schindelin et al., 2012; Tinevez et al., 2017). We then export the x-y-z-t coordinates of paired centrosomes into MATLAB, which permits rapid plotting and user-scoring of inter-centrosome distance (spindle length) versus time graphs and automated extraction of mitotic features. Manual track curation and pairing is time- and labor-intensive and large-scale analyses, such as genetic screens, are not realistic using this approach. Tracking is time-consuming largely because residual, global animal movement can mask individual centrosome movement, leading to a high frequency of tracking errors. Furthermore, centrosome pairing requires visual assessment and manual labelling of each putative pair. To address these challenges, we constructed CentTracker, a largely automated analysis pipeline, based around a series of integrated modules (Figure 3A).

**Figure 3.**
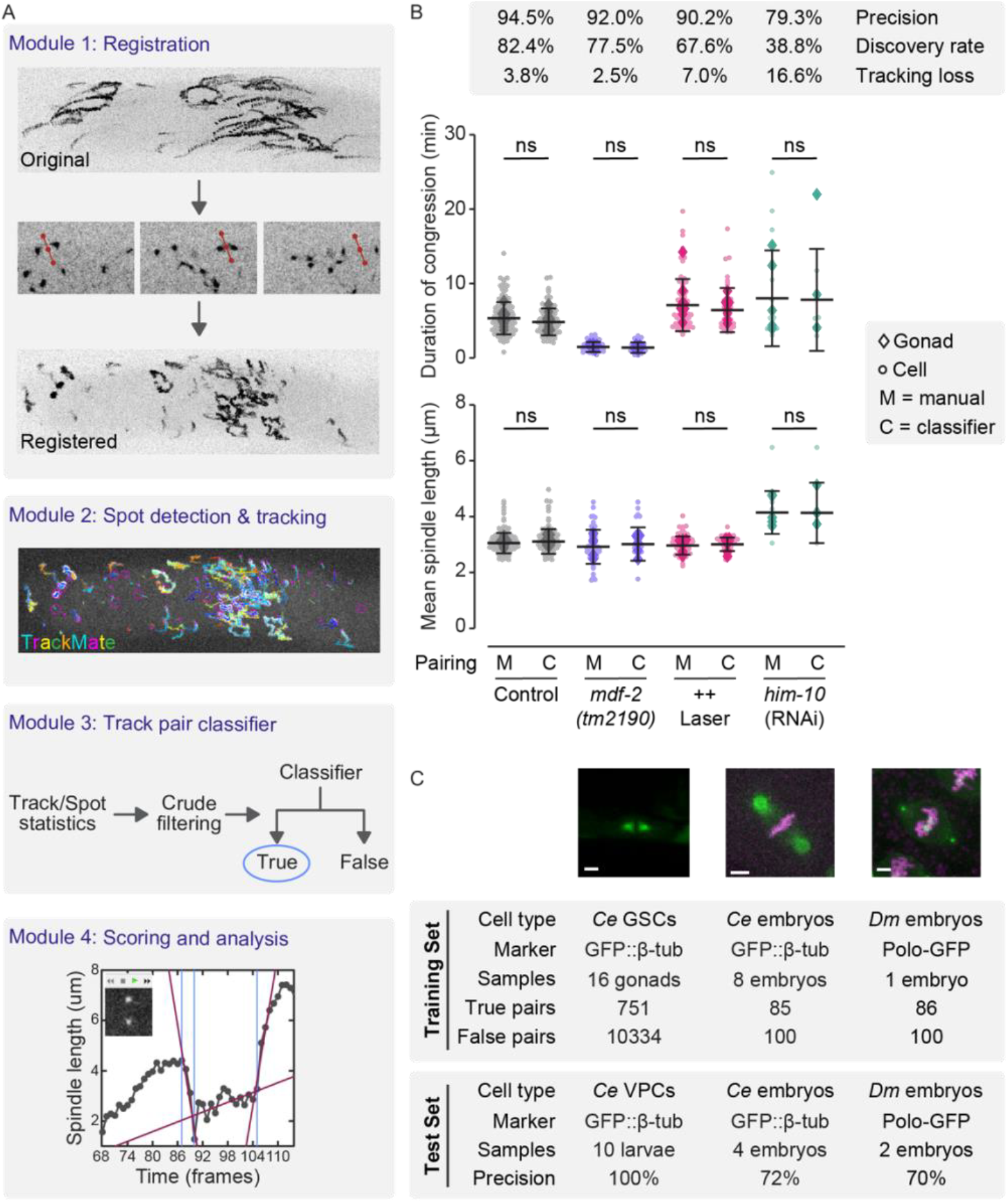
CentTracker enables automated monitoring of GSC mitosis. (A) Schematic overview of CentTracker workflow showing the tasks performed by each module. See main text for details. (B) Comparative analysis of the duration of congression (top) and mean spindle length (bottom) for centrosome pairs assigned manually (M) or using the CentTracker classifier (C), in phenotypically wildtype animals (grey; mCH::β-tubulin & mNG::ANI-1; n = 10 animals, manual ground truth n = 131 cells, classifier n = 108 cells) and in animals mutant for *mdf-2* (purple; *mdf-2(tm2190)*; GFP::β-tubulin; n = 6 animals, manual ground truth n = 40 cells, classifier n = 31 cells), imaged with 6-fold increase in laser intensity (magenta; mCH::β-tubulin & mNG::ANI-1; n = 9 animals, manual ground truth n = 71 cells, classifier n = 48 cells) or depleted for HIM-10/Nuf2 (green; GFP::β-tubulin; *him-10*(RNAi); n = 5 animals, manual ground truth n = 18 cells, classifier n = 7 cells). Circles represent individual cells. Diamonds represent the mean per animal. Black bars show the mean of the per cell values; error bars represent the standard deviation. Pairwise, manual versus classifier, comparisons were performed using a two-tailed Student’s *t*-test, except for *him-10*(RNAi), where a Wilcoxon Rank Sum test was used. ns = not significant (p ≤ 0.05). CentTracker’s precision (true positive rate), discovery rate (identified cells with complete congression as a percent of ground truth) and tracking loss (change in the number of identified cells with complete congression upon presenting the classifier with “error-free” tracks, as a percent of ground truth) for each condition is at the top. (C) Representative images (top) and summary tables for the training (middle) and performance (bottom) of CentTracker for pairing centrosomes in *C. elegans* (*Ce*) vulval precursor cells (VPCs, left), *C. elegans* embryonic blastomeres (center) and *Drosophila* (*Dm*) syncytial embryos (right), using the indicated marker to track centrosomes. Scale bars = 3 μm.

**Figure 4.**
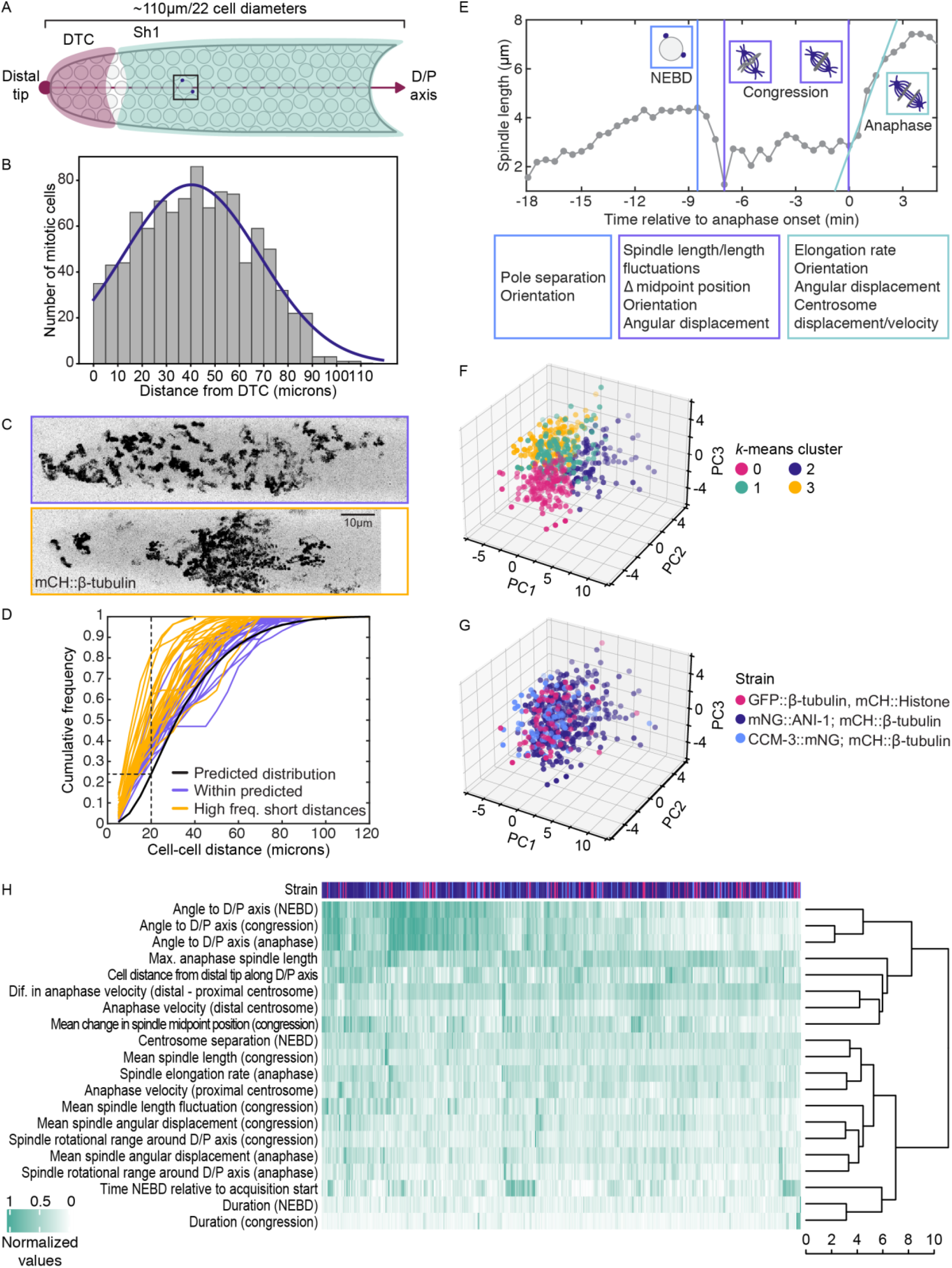
The spatial distribution of GSC mitoses is non-random, but mitotic features do not identify distinct sub-populations. (A) Schematic representation of the mitotic region of a late L4 gonad, which encompasses about 110 μm or ~22 cell diameters in length. Image registration in CentTracker allows for the distal tip (magenta circle) and distal-proximal (D/P) axis (magenta line; defined as a straight line through the center of the gonad arm from the distal tip towards the proximal end) to be marked in a single frame and propagated through all time points. The distal tip cell (DTC) is shown in magenta, the Sh1 sheath cell is shown in green. (B) Histogram showing the number of GSC mitoses along the D/P axis, relative to the distal tip, in 5 μm/~1 cell diameter bins. Grey bars represent bin counts. Purple line shows the normal distribution fit. Mean ± SD = 40.25 ± 28.07 μm. n = 74 animals, 996 cells. (C) Representative images of time- and z-maximum intensity projections of the mCH::β-tubulin channel for two gonads, one with a fairly uniform distribution of divisions (top) and one with one large cluster of divisions (bottom). Maximum intensity projections along the time axis are shown to highlight the position of divisions throughout the time-lapse acquisition. Scale bar = 10 μm. (D) The two-point correlation function for 46 gonads, each with ≥ 10 mitotic cells, is plotted as the cumulative frequency of mitotic cell-to-cell distances. The black line represents the mean predicted outcome, if cells were normally distributed along the D/P axis according to the gaussian fit in B. Distributions that fall within the predicted outcome (p > 0.05) are in purple and those that differ significantly from it (p ≤ 0.05; two-sample Kolmogorov-Smirnov test) are shown in yellow. Dashed lines highlight the increased frequency of short (≤ 20 μm) mitotic cell-to-cell distances in most gonads. (E) Representative spindle length versus time (relative to anaphase onset) plot for a single cell showing the various mitotic landmarks and indicating the types of mitotic features extracted for the different mitotic stages (NEBD, congression and anaphase). (F-G) The results of PCA using 34 mitotic features, followed by *k*-means clustering. A set of 547 GSCs are plotted along the first three principle components and are color-coded by cluster assignment (F) or by strain of origin (G). (H) Heat map showing the normalized values for the subset of mitotic features listed on the left, measured for dividing GSCs from each strain of origin (top, with color coding as in G), and ordered by hierarchical clustering, with the linkage between clusters indicated by the dendrogram to the right. Dendrogram branch length represents the Euclidean distance between clusters.

We reasoned that correcting for sample movement would permit the generation of sufficiently error-free tracks in TrackMate, such that automated pairing would be feasible. Thus the first CentTracker module was built to perform accurate image registration. Image registration, in this case, is a non-trivial problem due to the lack of fiduciary landmarks in our images and the fact that our labelled objects (centrosomes) are themselves highly dynamic. We based our registration approach on the observation that, whilst individual centrosomes are dynamic, the position of the metaphase plate (i.e. the spindle midpoint) is relatively stable with respect to the animal and/or germline. Moreover, the spindle midpoint is readily visible, and thus easily marked. As the majority of animal movement occurs within the x-y plane, spindle midpoint marking can be performed on 2D maximum intensity z-projections, which further simplifies this step. The x-y-t coordinates of spindle midpoints are then used to generate a translation matrix by calculating midpoint displacement between adjacent time points. Applying this translation matrix effectively corrected for animal movement over time and was sufficient to constrain centrosome trajectories to a significantly smaller range (Figure 3A). This reduction in global movement also enables the identification of relevant tissue landmarks at a single timepoint, which can then be accurately propagated throughout the image acquisition. For *C. elegans* GSCs, we specified the distal tip and the distal-proximal axis of each gonad arm, which allowed us to position cells along the length of the mitotic zone relative to the distal end or niche (see Figure 4A).

Registered images are passed to TrackMate for spot (i.e. centrosome) detection and tracking (CentTracker Module 2). While image registration enhanced tracking accuracy, low SNR can lead to imperfect spot detection, which, in turn, can lead to spurious and/or incomplete tracks. In addition, GSC mitoses are often clustered (see Figure 4C-D) such that two centrosomes from neighboring cells may be closer to one another than to their true pair, and spindle characteristics may vary, particularly under experimental conditions where spindle formation is perturbed, which can confound accurate pairing using hard filtering criteria. To address these issues, we built a trainable track pair classifier (CentTracker Module 3) which uses a modified random forest algorithm (Pedregosa et al., 2011) and a standard machine learning routine. Briefly, a data refining set of crude filters are applied to spot and track descriptors taken from TrackMate (or any other tracking software) and then an algorithm classifies each possible pair of centrosome tracks as “true” or “false” pairs, based on eleven numerical features. The user provides training material to this algorithm, which allows the classifier to refine the relative weight of each parameter and generate a model that can be applied to unseen data (see Supplemental Methods).

Once a set of paired tracks has been generated, the coordinates of paired centrosomes, plus any additional, user-defined landmarks (e.g. the position of the gonadal distal tip) are passed to MATLAB (CentTracker Module 4). After data import, spindle length versus time graphs are presented to the user, which allows for rapid scoring of mitotic events (e.g. NEBD and anaphase onset) by simply clicking on the plot at the relevant position in time. In addition, if a graph is ambiguous, the user can call a time-lapse image, cropped and centered on the centrosome pair in question, for visual confirmation (Figure 3A).

To test the performance of CentTracker, we applied it to a subset of our previous, manually-tracked data (Figure 3B; 10 animals, 205 true centrosome pairs, representing GSCs in various stages of mitosis, 131 of which undergo complete congression, i.e. in which both NEBD and anaphase onset could be accurately determined). CentTracker identified 164 centrosome pairs, 155 of which were validated as true pairs, for a precision of 94.5%. Notably, CentTracker identified 82.4% (108/131) of centrosome pairs from cells going through a complete congression (hereafter “discovery rate”). We next asked whether the performance of CentTracker could be improved further by removing residual tracking errors. To do so, we manually corrected for tracking errors in TrackMate, which were most commonly due to track truncations and/or breaks and spot misassignments, and supplied CentTracker with “error-free” tracked files. In this case, precision was unchanged (94.5%), but a larger fraction of cells with complete congression were returned (113/131, 86.25%). Thus, compared to labour-intensive manual tracking, CentTracker identifies the majority (155/205, 75.6%) of true centrosome pairs with high precision, including a proportional number of cells that undergo a complete congression (63.9% versus 69.6% of true pairs, manual versus CentTracker, respectively). Persistent tracking errors account for a tolerable loss (<4%) in discovery rate. Importantly, several features of mitosis, including duration of congression and spindle length, were similarly distributed when comparing cells identified using manual versus CentTracker pairing (Figure 3B), indicating that CentTracker is not biased for a certain subset of the population.

While CentTracker performed well on an unseen control data set, we wished to determine whether it would be robust enough to detect cells with aberrant mitoses. To this end, we challenged CentTracker with aberrantly short GSC mitoses (*mdf-2/Mad2(tm2190)* mutants, (Gerhold et al., 2015), delayed GSC mitoses (6-fold increase in laser intensity, this work), and severely perturbed spindle stability and delayed mitoses (*him-10/Nuf2*(RNAi) animals, (Gerhold et al., 2015); Figure 3B). CentTracker performed well for aberrantly short and delayed mitoses (precision: 92 and 90.2%; discovery rate: 77.5 and 67.6%, respectively), but struggled when faced with cells with severe spindle perturbations (*him-10/Nuf2*(RNAi); precision: 79.3%; discovery rate: 38.8; Figure 3B). Correcting for tracking errors prior to pairing lead to a moderate increase in the number of informative cells found in these animals (discovery rate = 55%, corresponding to a 16.6% tracking loss). In all cases, mitotic features (duration of congression and spindle length) were similarly distributed in datasets generated by CentTracker versus those generated manually (Figure 3B). Thus, even in animals with perturbed mitoses and fewer cells identified, CentTracker can produce a relatively unbiased sampling of the underlying population, which should be sufficient to detect interesting phenotypes.

Finally, to test the generalizability of the CentTracker software we applied it to several other cell types (Figure 3C). First, we used CentTracker to identify centrosome pairs in *C. elegans* larval vulval precursor cells (VPCs), in which centrosomes were also marked with GFP-tagged β-tubulin. In this case, we did not have a large enough dataset to train a new model; however, even using the classifier trained on mitotic GSCs, CentTracker was able to identify 55% of dividing VPCs with 100% precision. To test the “trainability” of CentTracker, we trained our classifier on *C. elegans* embryos, in which centrosomes were labelled with β-tubulin::GFP, and *Drosophila* embryos, in which centrosomes were marked with a GFP-tagged version of Polo kinase (Archambault et al., 2008; Moutinho-Santos et al., 1999). In both cases, even though the model was trained on a relatively small dataset (85/100 and 86/100 true/false pairs, respectively), the software identified true centrosome pairs in unseen data with 72% and 70% precision, respectively. Thus, CentTracker can be used to pair marked centrosomes in a range of mitotic cell types, even with sub-optimal centrosome markers, such as Polo kinase, which is also strongly recruited to kinetochores and the midbody, complicating tracking/pairing tasks (Archambault et al., 2008; Moutinho-Santos et al., 1999).

### GSC divisions are not randomly distributed along the length of the mitotic zone in L4 larvae

We next took advantage of CentTracker to collect data for a large population of mitotic GSCs (n = 996 cells; 74 animals, 3 genotypes/strains) and used this data set to ask how different aspects of mitosis were distributed across the GSC population. We first assessed the spatial distribution of GSC divisions within the mitotic zone (Figure 4A). We found that the frequency of divisions along the gonadal distal-proximal (D/P) axis followed a roughly normal distribution, with fewer divisions in the distal-most region, a peak in mitoses approximately 8 cell diameters (40μm) from the distal tip and very few divisions in the most proximal region of the mitotic zone (Figure 4B). Thus, the frequency of divisions in late L4 larval germlines follows a similar pattern to that which has been observed in adults (Crittenden et al., 2006; Gordon et al., 2020; Hansen et al., 2004; Maciejowski et al., 2006). Anecdotally, we observed that GSC divisions tended to be clustered together within the same germline. To determine whether our data supported this, we calculated the distribution of pair-wise distances between dividing cells, using the two-point correlation function (TPCF), in all germlines with at least 10 divisions (n = 46 animals; 821 cells; 3 genotypes/strains). We found that the majority of germlines (24/46) had a cumulative distribution of mitotic cell-to-cell distances that differed significantly from that predicted by our observed normal distribution of division frequency alone (Figure 4C-D). Further, most germlines (37/46) displayed a higher than expected frequency of short mitotic cell-to-cell distances (≤ 20μm or 4 cell diameters) and a subset of germlines showed a clear step-like TPCF, suggesting at least two discrete clusters, separated by a zone devoid of divisions (data not shown). Interestingly, analysis of fixed adult germlines has also suggested spatial clustering of GSC mitoses (Maciejowski et al., 2006), although the reasons for this are poorly understood. Thus, GSC mitoses are not randomly arrayed within the mitotic zone and show both a peak in frequency at intermediate distances from the distal tip and a tendency to occur within close proximity to one another.

### Mitotic properties do not identify discrete populations within the GSC pool

We next asked whether these spatial heterogeneities were accompanied by any population-level patterns in mitotic dynamics. We extracted a variety of mitotic features, including the duration of congression, centrosome positioning at NEBD, spindle length and length fluctuations, spindle rotation and orientation relative to the D/P axis, individual centrosome velocities, and spindle elongation rates (Figure 4A and E). To visualize cell-to-cell similarities in the GSC population, we performed Principal Component Analysis (PCA) across 34 defined mitotic features (Figure 4F). PCA is a common dimensionality reduction tool that identifies axes of maximal variance (principal components (PCs)) within multidimensional data. Plotting observations along these new axes can accentuate differences within the population and, when combined with a clustering algorithm (here *k*-means), identify subpopulations with a common collection of features. PCA was carried out on the subset of cells for which we were able to extract values for all mitotic parameters (n = 547). We found that less than 50% of the variance within our GSC population could be explained by the first three PCs (Figure S2A), and plotting all cells along these PC axes did not reveal any obvious subpopulations. *k*-means clustering analysis suggested an optimum cluster number of four; however, the predicted clusters are not well separated (Figure 4F and S2B). Together, these results suggest that GSCs do not form natural clusters based on the mitotic parameters extracted and that covariance between features is minimal. Importantly, neither genotype (Figure 4G), the time of mitotic entry relative to the start of imaging (Figure S2C), nor the position of a cell along the D/P axis (Figure S2D) were predictive of a cell’s position within PC space.

To investigate potential correlations between mitotic features while preserving feature identity, we generated heat maps for each feature and performed hierarchical clustering (Figure 4H and S2E). As expected, features derived from related measurements (e.g. centrosome displacement and velocity) clustered together (Figure S2E). Notably, none of our assayed features, many of which might be expected to inform on spindle assembly and/or stability, correlated appreciably with the duration of congression (Figure 4H). Thus, while variation in the duration of congression in GSCs is SAC-dependent (Gerhold et al., 2015), our analysis of mitotic features in GSCs undergoing unperturbed mitoses does not identify a specific aspect of spindle assembly that is strongly predictive of difficulties satisfying the SAC. Similarly, position along the D/P axis of the gonad did not correlate with any mitotic feature (Figure 4H), suggesting that spatial distinctions within the GSC population, which may potentially be related to “stemness” (Hubbard and Schedl, 2019), do not impact core mechanisms of mitotic progression.

### Spindle orientation in GSCs is biased towards the gonadal D/P axis in L4 larvae

The most strongly correlated features within our data related to spindle orientation along the D/P axis of the gonad. We found that a cell’s spindle orientation is similar throughout mitosis, from NEBD through anaphase, suggesting that spindles might be rotationally confined relative to the gonadal axis (Figure 4H). Regulated spindle orientation is an astral microtubule-dependent process that can shape tissues and impact stem cell fate (Kulukian and Fuchs, 2013; Noatynska et al., 2012; Yamashita et al., 2003). Spindle orientation in GSCs was proposed to be arbitrary with respect to the D/P axis (Crittenden et al., 2006; Kimble and Hirsh, 1979), suggesting that regulated asymmetric divisions, which generate two daughter cells that are intrinsically different from one another, do not contribute to stem cell maintenance and that GSCs are maintained as a population, with stem cell fate dictated by proximity to the niche (Morrison and Kimble, 2006). It has been shown recently, however, that spindle orientation in adult GSCs is biased at the interface of the DTC and proximal somatic sheath cell (Sh1), such that one daughter cell remains in contact with DTC (niche) appendages, while the other moves under the sheath cell and differentiates (Gordon et al., 2020), suggesting that a subset of GSC divisions may be asymmetric. In late L4 larvae, this interface is relatively smooth and orthogonal to the gonadal axis (Figure 4A; (Gordon et al., 2020), such that divisions biased along it will also be biased relative to the D/P axis.

To address whether spindle orientation in late L4 larvae showed any orientation biases, we measured the angle formed by each spindle and the gonadal D/P axis (Figure 4A). We assessed spindle orientation in 3D to capture the full range of spindle movements and to avoid potential pitfalls associated with 2D projected measurements (Jüschke et al., 2014). We compared our measurements to the theoretical distribution for random orientation along the gonadal axis, using a model described by Jüschke, et al., wherein the probability of a spindle assuming a particular orientation is proportional to the surface area of the sphere at this angle (Figure 5A; (Jüschke et al., 2014). Compared to this calculated random distribution, we found a strong enrichment of spindles that were oriented towards the D/P axis, and a pronounced lack of spindles in the orthogonal orientation (Figure 5B-C). As predicted by our hierarchical clustering analysis (Figure 4H), this orientation bias was observed throughout mitosis, i.e. at NEBD, during congression and in anaphase (Figure 5C).

**Figure 5.**
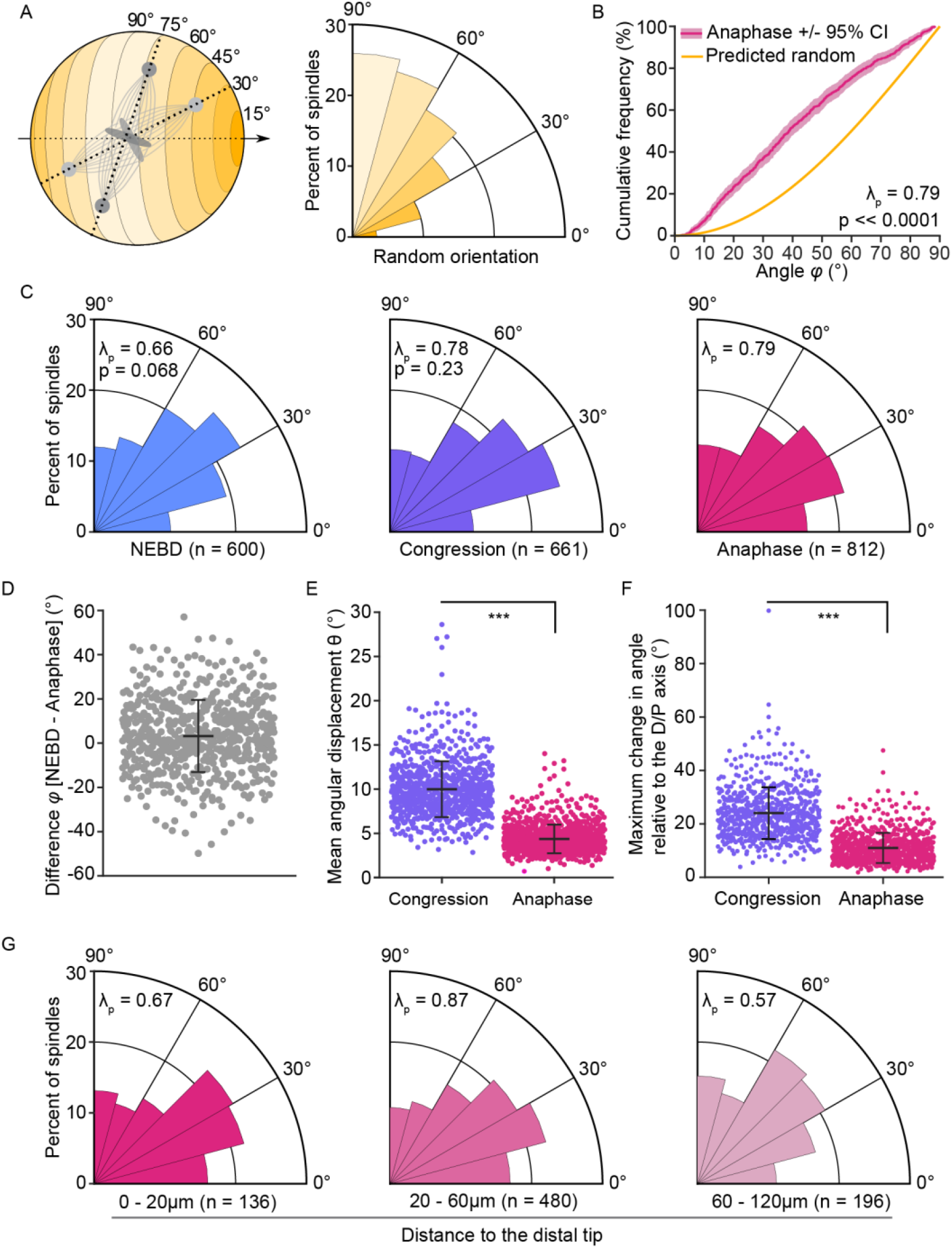
GSC spindles preferentially orient along the gonadal D/P axis in L4 larvae. (A) Schematic representation (left) of a cell modeled as a sphere, with the D/P axis (black, dashed, arrowed line) running horizontally, through the sphere’s center, from pole to pole. The cortical surface sampled by centrosomes from spindles oriented at different angles to the D/P axis are shaded according to the spindle angle formed, from parallel (0°; dark yellow), to orthogonal (90°; pale yellow). Two mitotic spindles are shown at angles of 30° (light grey) and 75° (dark grey) relative to the D/P axis. The polar histogram (right) represents the predicted distribution of spindle angles relative to the D/P axis, if orientation is random with respect to this axis. The probability in each bin is proportional to the surface area that can theoretically be sampled by centrosomes on spindles with a given orientation. (B) Cumulative distribution for spindle angles relative to the D/P axis for the calculated random distribution (yellow line) and the measured angles at anaphase (magenta line, mean with 95% confidence interval shaded, n = 812 spindles). The distribution at anaphase is significantly enriched for horizontal angles, i.e. towards the D/P axis, compared to random (p = 2×10^−61^, one-sample Kolmogorov-Smirnov test). (C) Polar histograms showing the distribution of spindle orientation relative to the D/P axis, at NEBD (blue, left) and during congression (purple, middle) and anaphase (magenta, right). (D) Beeswarm plot showing the difference (per cell) between the angle of the spindle relative to the D/P axis at NEBD versus during anaphase (mean ± SD = 3.2 ± 16.3°; n = 574 spindles). (E-F) Beeswarm plots showing the per cell, mean, spindle angular displacement (E) and the maximum change in mitotic spindle angle relative to the D/P axis (F) during congression (purple; mean ± SD = 10.0 ± 3.2° (E) and 24.0 ± 9.7° (F)) and anaphase (magenta; mean ± SD = 4.4 ± 1.6° (E) and 11.0 ± 5.7° (F)). The difference in values between congression and anaphase are significant (p < 1×10^−100^; paired-sample Student’s *t*-test). In D-F, black bars represent the mean and error bars show the standard deviation. (G) Polar histograms showing the distribution of spindle angles, relative to the D/P axis, binned by distances from the gonad’s distal tip. Bin edges and n are given underneath. In C and G, all distributions show a horizontal enrichment (λ_p_ > 0.50; i.e. towards the D/P axis) and all distributions are statistically different from random (p < 0.01; one-sample Kolmogorov-Smirnov test) but not from one another (p > 0.05; two-sample Kolmogorov-Smirnov test), except for the distribution of spindle angles within the most proximal bin (G, 60-120 μm) which is significantly different from the distribution in the middle, 20-60 μm, bin (p = 0.012; two-sample Kolmogorov-Smirnov test).

While we see a similar bias in spindle orientation at NEBD and anaphase at a population level, and a relatively strong correlation between these two values on a per cell basis (r = 0.75, p << 0.001), we found that spindles tend to skew towards the D/P axis as GSCs progress from NEBD to anaphase (mean = 3.2°; Figure 5D). Furthermore, the change in spindle angle between NEBD and anaphase varies widely (StDev of 16.3°) and can reach as high as 57° (Figure 5D). Thus while anaphase spindle orientation, relative to the D/P axis, is generally prefigured by centrosome positioning prior to NEBD, significant changes in spindle orientation can occur between these two events. The end result, however, is that spindles, on average, adopt an orientation more parallel to the D/P axis by anaphase.

Spindle movements (such as rotation) are typically achieved by cortical forces that are applied on astral microtubules and that can cause spindle oscillations, with an amplitude correlating with force magnitude (Grill and Hyman, 2005; Pecreaux et al., 2006). To determine if GSC spindles undergo oscillations, we measured the change in spindle angle between timepoints (angular displacement), during congression and anaphase. We found that angular displacement was significantly greater during congression than during anaphase (Figure 5E) and the maximum rotational range relative to the D/P axis was also much broader (Figure 5F). Thus, while the orientation of the mitotic spindle relative to the D/P axis varies during mitosis, it is largely set by the start of anaphase. Further, the fact that spindles tend to oscillate more around their midpoint during congression than in anaphase suggests that the loss of cohesion between sister chromatids contributes to stabilise the net force applied on astral microtubules during anaphase. Finally, the pair-wise oscillatory movement of the two centrosomes was nearly equal during congression (average difference in displacement between the two centrosomes: 0.1 ± 0.1 μm), which is different from what is observed in the asymmetrically dividing 1-cell *C. elegans* embryo, where force imbalance results in a significantly greater oscillatory movement of the posterior centrosome compared to the anterior one (Grill et al., 2003; Pécréaux et al., 2016). This suggests that forces are largely balanced on each side of the spindle during congression and is compatible with the notion that GSC divisions are symmetric.

Finally, we asked whether anaphase spindle orientation, which is predictive of daughter cell positioning after cytokinesis, varied along the length of the gonad. To test this, we used our measured distribution of mitotic entries along the mitotic zone and published reports on the position of the DTC and the DTC-Sh1 interface in late L4 larvae to define three bins: (1) GSCs mainly dividing under the DTC cap (0-20μm, covering ~4 cell diameters from the distal tip; (Crittenden et al., 2006), GSCs near the DTC-Sh1 interface (20-60 μm, comprising the zone of peak GSC mitosis, which we see at ~40μm or 8 cell diameters; (Gordon et al., 2020), and GSCs dividing in a region covered by Sh1 only (60-120 μm, extending to the transition zone, ~22-24 cell diameters from the distal tip; (Crittenden et al., 2006; Gordon et al., 2020). We found that anaphase spindle orientation in each of these three regions is statistically different from the calculated random distribution and is significantly more biased towards the D/P axis (Figure 5G), although this bias is less pronounced for spindles in GSCs within the Sh1-only, most proximal regions of the mitotic zone. Thus, unlike in adults, spindle orientation in GSCs in late L4 larvae is generally oriented toward the D/P axis in all regions of the gonad, and does not appear to be strongly influenced by the DTC-Sh1 interface. Altogether our results suggest that while larval GSCs tend to divide along the D/P axis, this bias in orientation is not accompanied by evidence of regulated asymmetric cell division, neither in terms of spindle dynamics nor in terms of its relationship to relevant landmarks. One possibility is that underlying anisotropies in cell and/or tissue shape contribute to GSC spindle orientation, but further study is needed to address this.

In sum, this work provides the first systematic analysis of technical factors that affect the division of *C. elegans* GSCs in live imaging experiments. While animal starvation is the main technical factor impacting GSC mitosis, our work demonstrates that its effects are minimal within the first 40 minutes of acquisition, providing a window during which GSCs division can be visualized under seemingly physiological conditions. We further introduce CentTracker as a flexible image analysis pipeline, which facilitates the extraction of GSC mitotic features from large-scale imaging data sets and which is adaptable for analyses of other cell types and other organisms. Finally, we use CentTracker to analyse several features of GSC mitosis, providing evidence for spatial clustering of GSC divisions and a penetrant bias in spindle orientation towards the D/P axis of the gonad.

## Materials and methods

### *C. elegans* strains and culture

*C. elegans* animals were maintained at 20°C on nematode growth medium (NGM) and fed with *Escherichia coli* strain OP50 according to standard protocols (Brenner, 1974). Synchronized L1 larvae were obtained by collecting gravid hermaphrodites and dissolving them in a solution of 1.2% sodium hypochlorite and 250 mM sodium hydroxide. Embryos were collected, washed in M9 buffer (22.04 mM KH_2_PO_4_, 42.27 mM Na_2_HPO_4_, 85.55 mM NaCl, 1 mM MgSO_4_) and hatched for 24h at 15°C in M9 buffer. Late L4 larvae were obtained by inoculation of synchronized L1 larvae on NGM plates containing 1 mM isopropyl β-d-thiogalactoside (IPTG) and 25 μg/ml carbenicillin (Carb) and fed for 44-48 hours with *E. coli* strain HT115 transformed with the empty RNAi control vector L4440 (from the Ahringer RNAi collection; (Kamath et al., 2003)). Consistent with previous reports (Chaudhari et al., 2016), we find that animals reared on HT115 bacteria have a more uniformly high frequency of GSC divisions (data not shown).

Strains imaged for this study were UM679: *ltSi567[pOD1517/pSW222; Pmex-5::mCherry::tbb-2::tbb-2_3’UTR; cb-unc-119(+)]I; ani-1(mon7[mNeonGreen^3xFlag::ani-1]) III*, ARG16: *ltSi567[pOD1517/pSW222; Pmex-5::mCherry::tbb-2::tbb-2_3’UTR; cb-unc-119(+)]I; ccm-3(mon9[ccm-3::mNeonGreen^3xFlag]) II; unc-119(ed3) III*, and JDU19: *ijmSi7 [pJD348/pSW077; mosI_5’mex-5_GFP::tbb-2; mCherry::his-11; cb-unc-119(+)]I; unc-119(ed3) III*.

### Worm mounting and live imaging

Animals were anaesthetized in M9 buffer containing 0.04% tetramisole (Sigma) and transferred to a 3% agarose pad, molded with grooves made by a custom microfabricated silica plate, onto which a coverslip was placed, as described (Gerhold et al., 2015). The chamber was backfilled with M9 buffer containing 0.04% tetramisole and sealed using VaLaP (1:1:1 Vaseline, lanolin, and paraffin). Images (excepting those in Figure S1A, see supplemental methods) were acquired at room temperature (∼20°C) with an AxioCam 506 Mono camera (Zeiss) mounted on an inverted Cell Observer spinning-disk confocal microscope (Zeiss; Yokogawa), using a 60x Plan Apochromat DIC (UV) VIS-IR oil immersion objective (Zeiss), controlled by Zen software (Zeiss). Acquisitions used 200ms exposure with 488nm (35mw) and 561nm (50mw) solid-state lasers, both set at 10% of intensity. Confocal sections of 0.5μm were acquired over 38μm of depth, at 30 second intervals, for durations of 40 or 90 minutes.

### Survival and brood size assays

Imaged animals were recovered by gently lifting the coverslip and removing the animal by mouth pipet. Animals were washed several times in M9 buffer, before being transferred to individual seeded NGM plates. Survival was assessed by monitoring animal movement a few hours after the transfer. Animals were transferred to fresh plates every 24 hours for a total of 72 hours and brood size was determined by counting the total number of progeny present on each plate 48 hours after the animal had been removed.

### Manual GSC centrosome tracking and pairing

Centrosome identification and tracking were performed using the Laplacian of the Gaussian (LoG) blob detector and Linear Assignment Problem (LAP) spot linker in the Fiji (Schindelin et al., 2012) plugin TrackMate (Tinevez et al., 2017). Tracks were manually curated to correct for tracking errors and all spots within a given centrosome track were renamed to indicate association between appropriate pairs (centrosomes within the same cell; e.g. Cent1a and Cent1b).

### Scoring GSC mitotic events from pairs of tracked centrosomes

Centrosome x-y-z-t coordinates and labels were exported and analyzed using a custom MATLAB script. Centrosome-to-centrosome distance (spindle length) was plotted relative to time (frames) and graphs were used to define three mitotic events: nuclear envelope breakdown (the last frame prior to the rapid decrease in spindle length, which is coincident with the appearance of microtubules within the nuclear space and thus NEBD), the start of chromosome congression (the first frame, post-NEBD, at which spindle length reaches a stable minimum) and anaphase onset (the last frame prior to rapid spindle elongation). Mitotic events were scored manually, by clicking on spindle length versus frame plots, with the option of calling a cropped time lapse image of the cell in question for visual confirmation. The duration of congression was calculated from the points of intersection of three least-squares lines fit to (1) spindle length from four frames prior to and including the start of congression, (2) all frames between the start of congression and anaphase onset, and (3) four frames, including the end of congression and the three following frames. All subsequent data analysis was performed using these mitotic events as landmarks to align cells and calculate various mitotic features.

### CentTracker workflow

The workflow for CentTracker involves four steps: registration, tracking, track pairing and cell scoring (Figure 3A). To correct for sample movement in the x-y plane (registration; Module 1), users mark the spindle midpoint of single cell at each time frame, in a z-maximum projection of the time lapse image, using a set of ImageJ macros, which automate all steps aside from spindle midpoint marking. An x-y-t translation matrix for registration is constructed by calculating a 2D direction vector for each timepoint *v_t_* = *p_t_*_−1_ − *p_t_*, where p_t_ and *p_t_*_−1_ are midpoint positions of the same spindle at timepoints *t* and *t*-1, respectively. To combine translation matrices from different spindles, matrices are concatenated by time axis. Input movies are then registered by shifting all z-slices of time *t* by the direction vector *v_t_* and zero-padding the margins. Translation matrix generation and image registration are performed in Python. Spot detection and track construction (Module 2) are performed by TrackMate (Tinevez et al., 2017), as in the manual method (described above), but excluding the spot labelling and track curation steps. We note, however, that any spot detection and tracking paradigm could be used, as long as the output data is compatible with the track pairer (Module 3). Track pairing (Module 3) is performed using a trainable, machine-learning based approach, which is implemented in Python using scikit-learn (Pedregosa et al., 2011). Crude filters (the minimum duration of individual track pairs, the minimum number of frames in which two tracks co-appear, and the maximum distance between two tracks at any frame) are applied to the data and the resulting, curated data set is converted into a set of eleven nine numerical features, which are fed into a random forest classifier to generate true/false predictions. Model details are provided in the supplemental methods. The x-y-z-t coordinates of paired centrosomes are passed to MATLAB and analysed as for manual pairs, using custom scripts (described above). Step-by-step instructions, examples and software code/scripts are available to download in a GitHub repository (https://github.com/yifnzhao/CentTracker). The main modules of the CentTracker were developed using python 3.8.5. Existing packages and software are used whenever possible. The essential python libraries used in the CentTracker software include numpy (van der Walt et al., 2011), pandas (McKinney, 2011), scikit-learn (Pedregosa et al., 2011), and scikit-image (van der Walt et al., 2014).

### CentTracker datasets

The control dataset consists of GSCs from a total of 16 individuals from stains UM679 and JDU19, imaged for 40 or 90 minutes, for this manuscript. We used the control dataset to train, validate and evaluate the random forest models in CentTracker. We then evaluated the generalization ability of CentTracker on three unseen *C. elegans* GSC datasets with different degrees of perturbations: we used previously published image data from our labs for GSCs in *mdf-2*(*tm2190*) and *him-10*(RNAi) animals (Gerhold et al., 2015) and data from control animals imaged with a 6-fold increase in laser intensity (this work). We further applied CentTracker on 3 additional previously acquired, image datasets from different tissues and/or model systems: *C. elegans* vulva precursor cells and embryos (A. R. Gerhold, unpublished data), and *Drosophila melanogaster embryos* (V. Archambault, unpublished data).

### Definition of the distal-proximal (D/P) gonadal axis

The distal tip (D) was defined in 3D as the distal most point of the gonad, at the center-most z-slice of the image (i.e. the xy-plane which passes through the middle of the rachis for the majority of the mitotic zone). The D/P axis (α) was then defined as a straight line from the distal tip (D), along the middle of the tissue (rachis) toward the proximal region (P). The vector *α* representing the distal-proximal axis was assumed to be constant across timepoints and was calculated as:

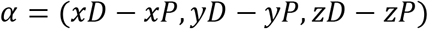

Animals with distorted (e.g. curved) sections of the gonad were excluded.

### Spindle orientation measurements in 3D

Spindle orientation calculations were carried out in MATLAB, unless otherwise noted. The spindle was represented as a vector *c* connecting the 3D coordinates (x, y, z) of the two centrosomes (C1, C2) at each timepoint of a registered movie:

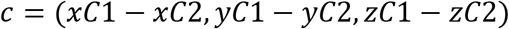

The orientation of the spindle relative to the D/P axis was defined as the angle *ϕ* measured between the vector *c* the vector *α,* representing the distal-proximal axis of the gonad (as described above). *Φ* was calculated at each timepoint using the following scalar product, where II II represents the vector’s norm:

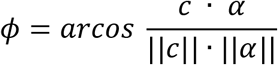

The change in mitotic spindle orientation between timepoints (angular displacement) was defined as the angle *ϴ* measured between the vector c_t_ connecting the two centrosomes at a given timepoint (C1_t1_, C2_t1_) and that connecting the same centrosomes at the next timepoint (c_t+1_). It was calculated at each timepoint using the following scalar product, where II II represents the vector’s norm:

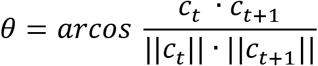

The theoretical random distribution of spindle angles to the D/P axis was calculated by adapting a method described previously (Jüschke et al., 2014), in which the probability of spindle poles orienting at a given angle is proportional to the surface area at this angle. A GSC was approximated as a sphere of radius 1 that is traversed in its center by a straight line defining the distal-proximal (0°) axis. The angle *ϕ* formed by the intersection of the spindle and D/P axes defined a cortical circumference (C):

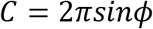

The surface area of the sphere *Aϕ* found between the D/P axis (0°) and the cortical circumference of a given angle (*ϕ*) and its symmetric counterpart (*-ϕ*, where 0° ≤ *ϕ* ≤ 90°) can be calculated as:

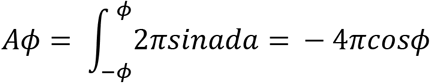

The fraction *fϕ* of the cortical surface area within this angle (*Aϕ*) to the total surface area of the sphere (A90° = 4π) can be determined as:

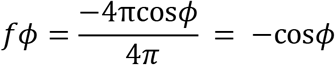

If mitotic spindle orientation were randomly distributed between parallel (0°) and perpendicular (90°) to the D/P axis, the probability *P* of a spindle orientation found between two given angles (*ϕ*_1_ and *ϕ*_2_, where 0° ≤ *ϕ*1 < *ϕ*2 ≤ 90°) can be calculated as:

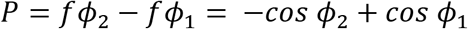

The deviation of measured GSC spindle orientation from the calculated theoretical random distribution in the parallel axis (λ_p_) was determined using the R fit parameter script, as previously described (Jüschke et al., 2014). The hypothesis that GSC spindle orientation is random was tested against the calculated theoretical random distribution by a one-sample Kolmogorov-Smirnov test (MATLAB kstest). To test the hypotheses that GSC spindle orientation is the same with respect to the mitotic stage and the position along the D/P axis, we applied two-sample Kolmogorov-Smirnov tests (MATLAB kstest2) with Bonferroni corrections.

### Spatial analysis of GSC divisions

To measure the frequency of mitotic GSCs as a function of distance along the D/P axis, the Euclidean distance in x for the mean spindle midpoint of each cell relative to the distal tip was calculated and the distribution was sorted into 24 bins, in Python 3.8 using the JupyterLab environment and key libraries including numpy 1.18.5 (Harris et al., 2020), pandas 1.0.5 (McKinney, 2011) and matplotlib 3.2.2 (Hunter, 2007). A gaussian fit to the distribution was determined using a Levenberg-Marquardt algorithm and least squares statistic, using the astropy package 4.0.1post1 (The Astropy Collaboration 2018, (Robitaille et al., 2013). The two-point correlation function (TPCF) for each gonad was computed in MATLAB by calculating the Euclidean distance in x-y-z between each cell and every other cell in the same gonad, using the 3D coordinates of the spindle midpoint at anaphase onset as the cell’s centroid. The resulting distances were binned by 5μm, and bin counts were converted to a cumulative frequency distribution. To calculate the predicted TPCF, a random gaussian distribution of 10,000 x coordinates, with parameters equal to the gaussian fit for measured mitotic frequency along the D/P axis, and a random uniform distribution of 10,000 y and 10,000 z coordinates, confined within the maximum range for these values for all gonads, were generated. For each gonad, 10,000 TPCF calculations with *n* cells, where *n* = the number of measured mitotic cells, were performed, using random draws with replacement from simulated x-y-z coordinates to create mitotic cell centroids. The average of these 10,000 iterations was taken as the representative predicted TPCF for a given gonad and was compared to the measured TPCF using a two-sample Kolmogorov-Smirnov test (MATLAB kstest2). Gonads enriched for cell-cell distances ≤ 20μm were defined as those for which the measured frequency was > twice the standard deviation for the simulated TPCF set.

### Large-scale analysis of GSC mitotic features

Mitotic features were calculated and feature lists were compiled in MATLAB. PCA and *k*-means clustering were performed in Python using scikit-learn 0.22.1 (Pedregosa et al., 2011), with the following modules: Data were standardized using sklearn.preprocessing.StandardScaler().fit_transform(); PCA was carried out using sklearn.decomposition.PCA(*n_components=34*, *svd_solver=‘auto’); k*-means clustering was carried out using sklearn.cluster.KMeans(n_clusters=4); Silhouette coefficient calculations and plots were performed using sklearn.metrics.silhouette_samples and sklearn.metrics.silhouette_score. Hierarchical clustering and heatmap generation were carried out using the R package ComplexHeatmap 2.4.3 (Gu et al., 2016), using the complete linkage clustering method with Euclidean as the distance metric.

### Additional statistical analysis

Curve fitting in Figure 1C was performed using the MATLAB Curve Fitting Toolbox. Comparison between two independent samples was performed using a two-tailed Student’s t-test (MATLAB ttest2) except for Figure 3B, *him-10*(RNAi), where a Wilcoxon rank sum test (MATLAB ranksum) was performed. Comparison of multiple means was performed using a Kruskal-Wallis test with a Tukey-Kramer post hoc test (MATLAB kruskalwallis and multcompare). Increased variability in the duration of congression (Figure 2C) was assessed by performing a Kruskal-Wallis test with a Tukey-Kramer post hoc test (MATLAB kruskalwallis and multcompare) on the distribution of sample residuals. If p < 0.05, we concluded that a treatment led to a more variable outcome.

## Acknowledgements

We are grateful to Dr. Vincent Archambault (IRIC, UdeM) for sharing unpublished movies of Drosophila syncytial embryos undergoing mitosis, as well as Drs. Amy Maddox (UNC Chapel Hill), and Arshad Desai (UC, San Diego) for sharing strains and reagents. We also thank Christian Charboneau of IRIC’s Bio-imaging facility and the staff at McGill’s Advanced BioImaging Facility (ABIF) for technical assistance, and members of the Hickson, FitzHarris, Labbé and Gerhold labs for help and advice. We thank Preshanth Jagannathan for advice on spatial clustering analysis. MRZ held an Alexander Graham Bell Canada Graduate Scholarship from the Natural Sciences and Engineering Research Council of Canada (NSERC), an IRIC Doctoral Scholarship and scholarships from UdeM’s molecular biology programs and the school of graduate and postdoctoral studies. YZ was partially supported by the Sheila Ann MacInnis Grant Science Undergraduate Research Award (SURA). This work was funded by a grant from the Canadian Institutes of Health Research (MOP-115171) to JCL and ARG. IRIC is supported in part by the Canadian Center for Excellence in Commercialization and Research, the Canada Foundation for Innovation and the Fonds de Recherche du Québec – Santé.

## Supplemental Figures, Figure Legends, Tables and Methods

**Figure S1.**
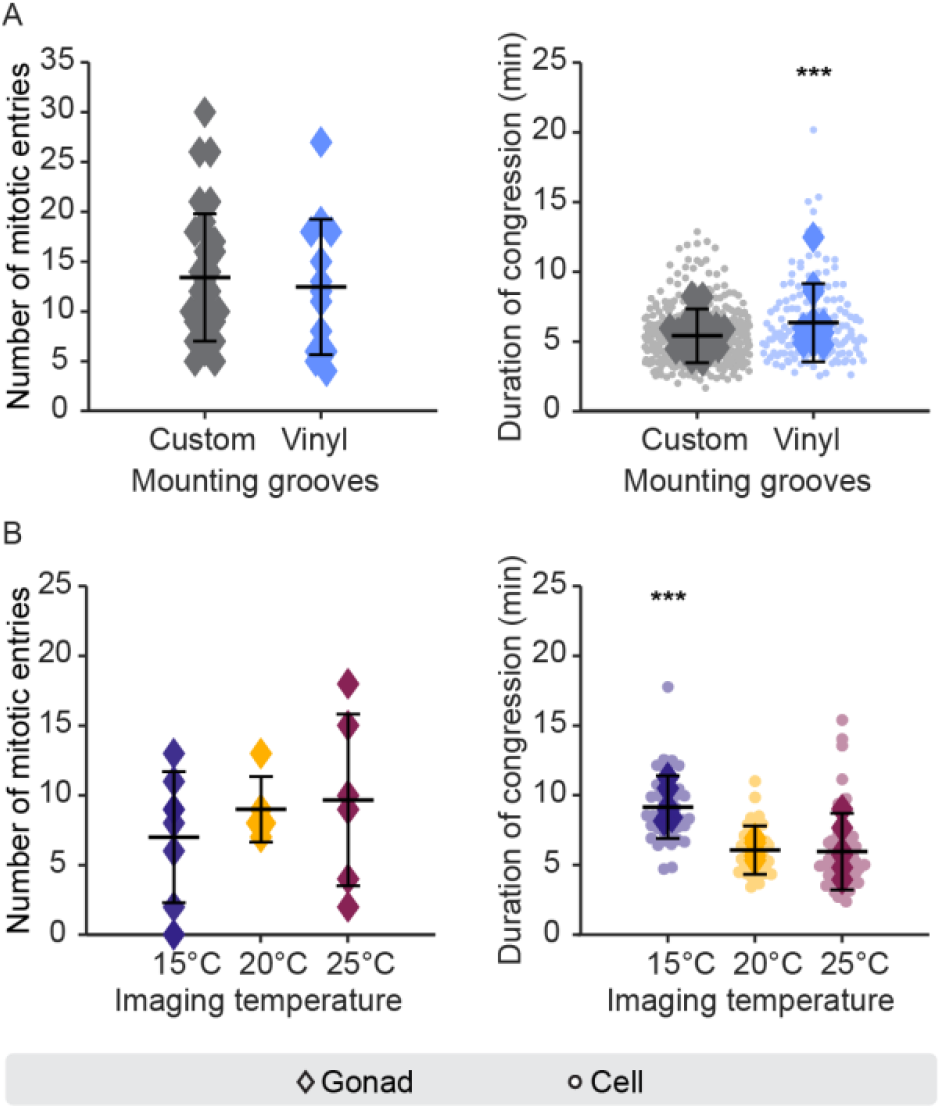
The effect of imaging temperature and using mounting pads molded with a vinyl LP record on GSC mitosis. (A-B) Beeswarm plots showing the number of mitotic entries per gonad (left) and the per cell (circles) and per gonad mean (diamonds) duration of congression (right) in (A) animals mounted on 3% agarose pads molded with grooves from a vinyl LP record (blue; n = 13 animals, 154 cells) compared to animals mounted under standard conditions and in (B) in animals raised at 20°C and imaged at 15°C (purple; n = 7 animals, 47 cells), 20°C (yellow; n = 5 animals, 37 cells) and 25°C (magenta, n=6 animals, 53 cells). Control data in A (grey) is reproduced from Figure 2. For A and B, black bars represent the mean of the per animal (left) and per cell (right) values; error bars represent the standard deviation. Statistical analyses were done using a two-tailed Student’s *t*-test (A) or the Kruskal-Wallis test with a Tukey-Kramer post hoc test (B). *** = p < 0.001.

**Figure S2.**
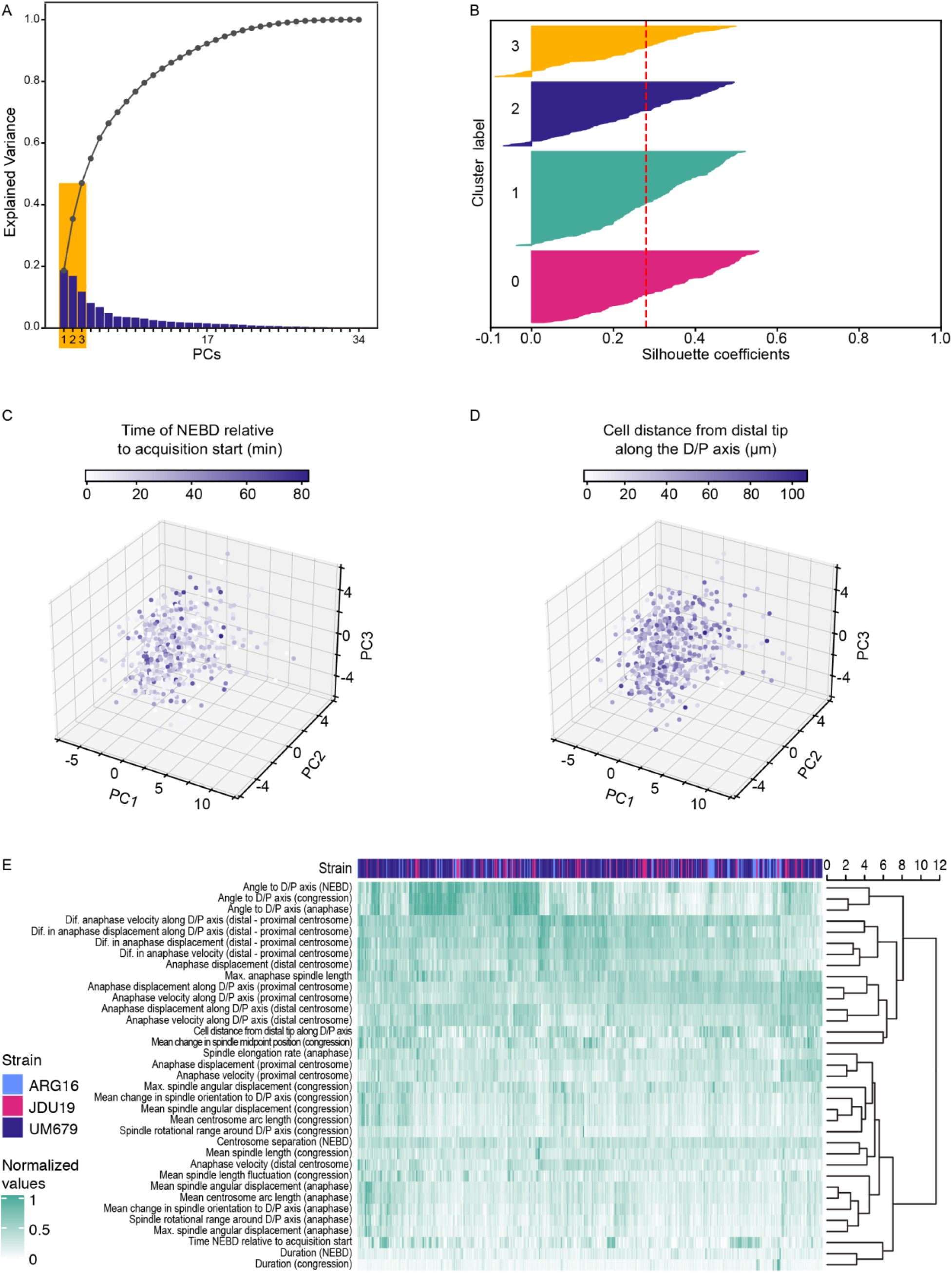
PCA and hierarchical clustering of GSC mitotic features does not reveal natural clustering. (A) Bar graph combined with cumulative distribution showing the percent of variance explained by each of the possible 34 principle components (PCs). The first three PCs (boxed in yellow) explain < 50% of the variance. (B) Plot showing the silhouette coefficients for all observations (here each a GSC) sorted by cluster, as determined by *k*-means clustering. The silhouette coefficient represents how similar an observation is to its own cluster, relative to the nearest neighboring cluster, with similarity calculated using Euclidean distance, and is a measurement of how well a given observation fits within its assigned cluster. The mean silhouette score is shown in red. (C-D) PCA plots for 547 GSCs plotted along the first 3 PCs and colored-coded according to their time of mitotic entry (NEBD) relative to the start of image acquisition (C) or their position in the mitotic zone relative to the distal tip of the gonad (D). (E) Heat map showing the normalized values for all 34 mitotic features (listed on the left), for GSCs from each strain of origin (top), and ordered by hierarchical clustering, with the linkage between clusters indicated by the dendrogram to the right. Dendrogram branch length represents the Euclidean distance between clusters.

**Table S1.**
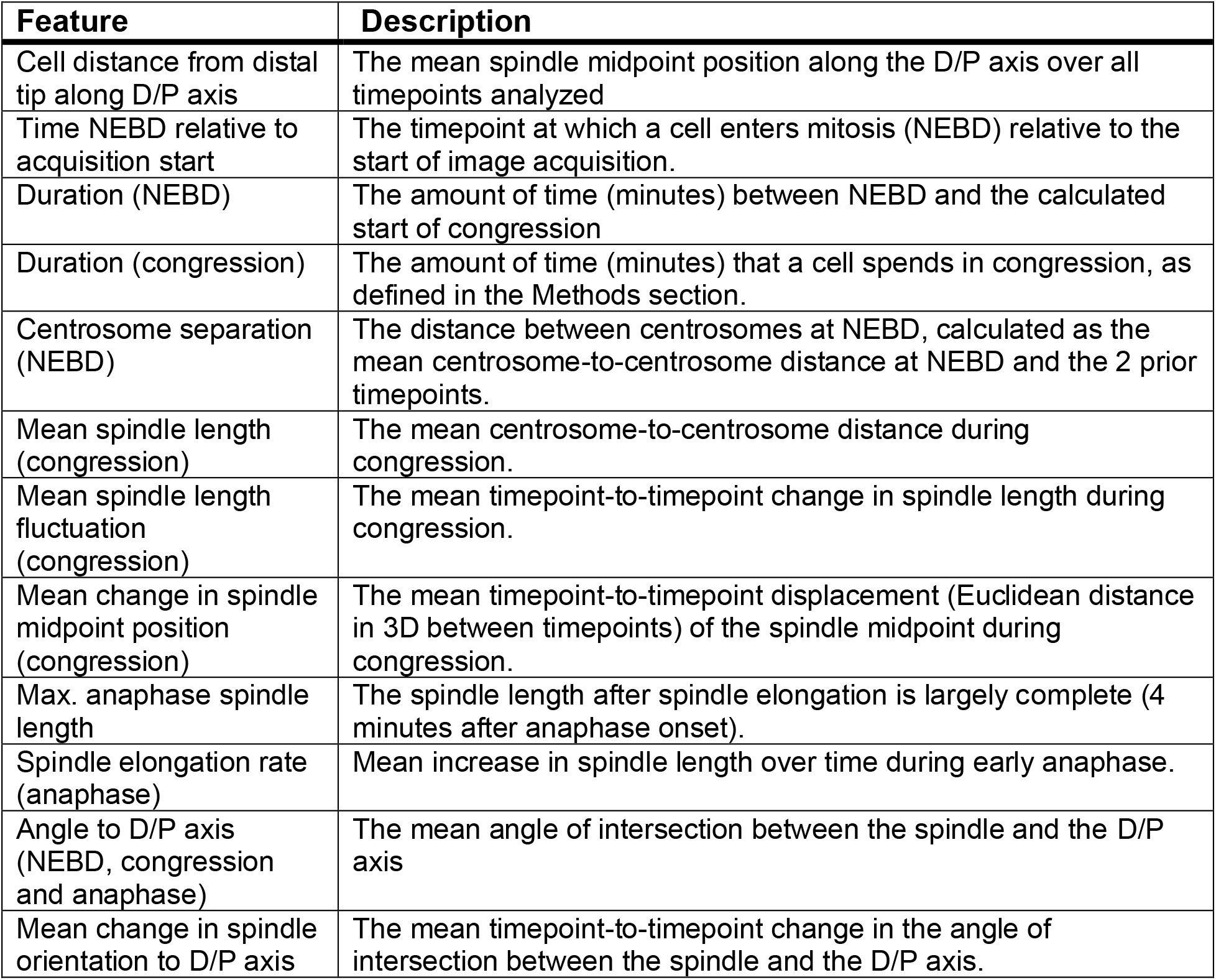
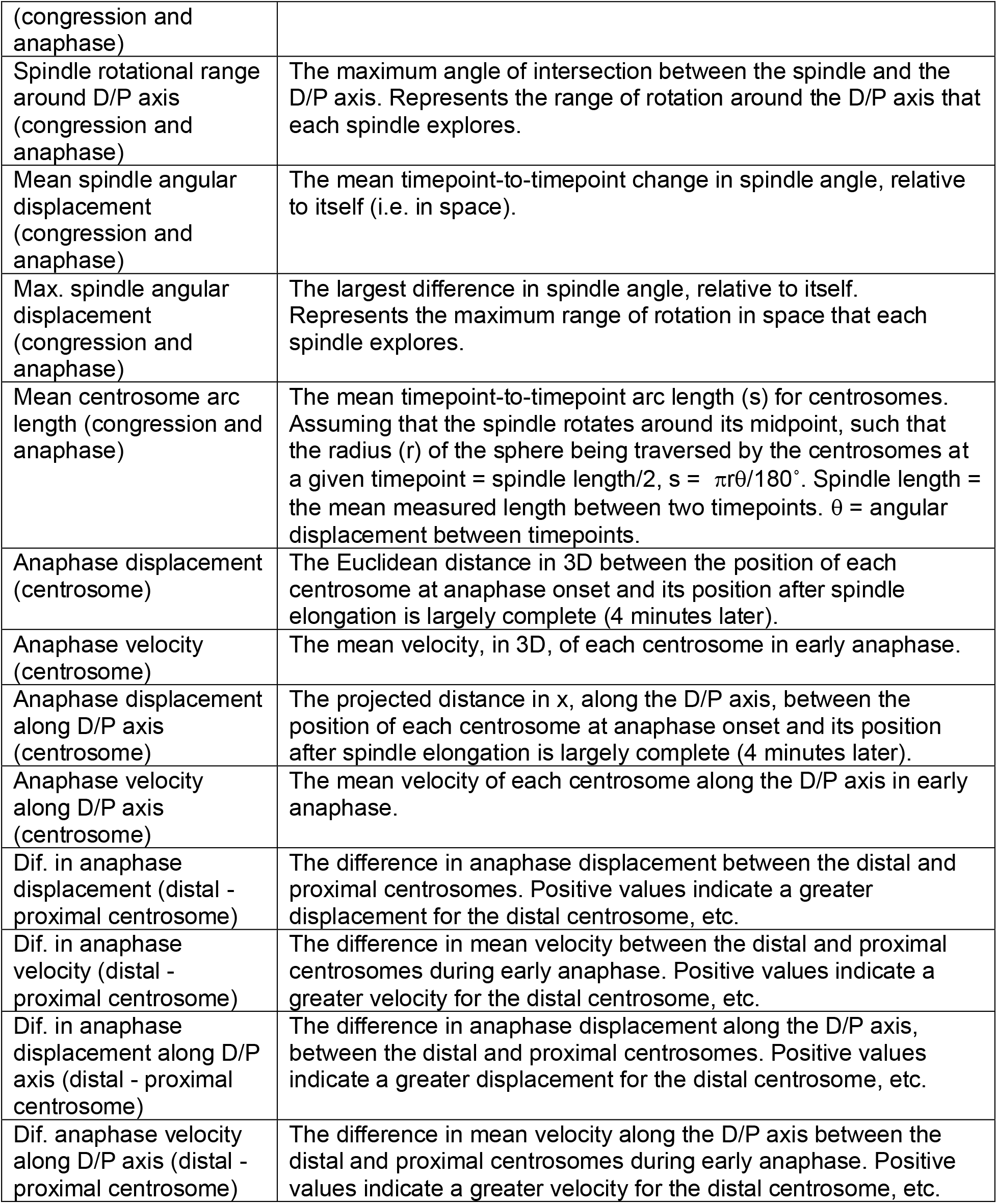
Descriptions of the mitotic features extracted from centrosome tracking and spindle length measurements in GSCs. Features were measured for NEBD, congression and anaphase as follows: NEBD values are calculated over 3 timepoints, the user-defined NEBD and the two preceding timepoints, except for the duration of NEBD, which is the amount of time between the timepoint marked as NEBD and the calculated start of congression. Congression values are calculated over all timepoints within the congression period. Anaphase values fall into two categories: *early anaphase* values are calculated over 5 timepoints, the last timepoint of congression and the following 4 timepoints (2 minutes), which correspond to the period of rapid spindle elongation; *anaphase* values are calculated over 9 timepoints, the last time point of congression and the following 8 timepoints (4 minutes), at which point spindle elongation is largely complete. Where defined, the distal and proximal centrosomes are the centrosome closest to and farthest from the distal tip of the gonad, respectively.

## Supplemental Methods

### Worm mounting and live imaging for Figure S1B

Animals were mounted as described in Methods. Images were acquired on a Quorum WaveFX-X1 spinning disk confocal, controlled by MetaMorph software, using a Leica 63x/1.40-0.60** oil HCX PL APO objective and 50mW 491nm and 568nm diode lasers. Confocal sections of 0.75μm were acquired over 19.5μm for a duration of 40 minutes, using dual camera mode, with ET 525/50 and FF 593/40 emission filters, 200ms exposure time and two Photometrics PRIME BSI CMOS cameras. Sample temperature was regulated using a CherryTemp microfluidic temperature control device (Cherry Biotech; (Velve Casquillas et al., 2011)).

### CentTracker track pair classifier (TPC) construction and validation

#### 1.1 TPC algorithm overview

We applied the random forest algorithm (Breiman, 2001; Louppe, 2015), a renowned machine learning classification technique for the track pair classification task. In this study, we adopted the random forest implemented by the machine learning platform scikit-learn (Pedregosa et al., 2011), where the final classification is computed by averaging the probabilistic predictions of all tree predictors, as opposed to the majority voting in the original method (Breiman, 2001). The final predicted outcome can be interpreted as the conditional probability of the class given the input.

#### 1.2 TPC feature construction

For each putative track pair, we constructed the following nine features as input to TPC:

1. The initial spindle length, i.e., the distance (“spindle length”) between two tracks when they first co-appear;
2. The final spindle length, i.e., the distance (“spindle length”) between two tracks when they last co-appear;
3. The maximum spindle length, i.e., the maximum distance between two tracks at any time frame;
4. The minimum spindle length, i.e., the minimum distance between two tracks at any time frame;
5. The track duration, i.e., the duration in which both tracks are present;
6. The congression duration, approximated by the number of continuous time points in which the spindle length is under 4 microns (by default);
7. The center standard deviation, i.e., the standard deviation of the midpoint of two tracks at all time points when the tracks are both present;
8. The normal standard deviation, where the normal is calculated by taking the vector difference between the track i and track j (the order here is determined arbitrarily but remain the same for the given two tracks);
9. The max intensity, i.e., the maximum of sum of all the values for pixels within the physical radius from the spot center in all time frames;
10. The contrast, defined as:

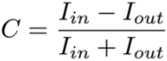

where *I_in_* is the mean intensity inside the spot volume (using the physical radius), and *I_out_* is the mean intensity in a ring ranging from its radius to twice its radius.
11. The average estimated diameter of two tracks from the TrackMate output, based on contrast calculation.

It should be noted that, after the calculation of the numerical features for all of the putative pairs, we applied max-min normalization to the max intensity and contrast terms, per movie, to adjust for the movie-level intensity differences.

#### 1.3 TPC experimental settings

We followed a standard machine learning routine to train, validate and evaluate the classifier. The data was split into a training (66.6%) set and a test set (33.3%). To address the imbalanced class distribution present in the dataset (false pairs are approximately 9 times more prevalent than true pairs), we applied a stratified 3-fold cross-validation during the hyper-parameter tuning to represent the original class distributions across each train-validation fold. The model with the highest accuracy at the hyper-parameter tuning is selected as the final model.

We used three metrics to evaluate the model prediction performance, including accuracy, F1 score and average precision. Accuracy score is defined as:

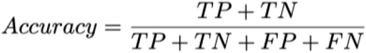

where *TP*, *TN*, *FP,* and *FN* stand for true positive, true negative, false positive, false negative, respectively. F1 score is the harmonic mean of the precision P and recall R, where an F1 score reaches its best value at 1 and worst score at 0, defined as:

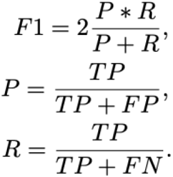

The average precision (AP) score summarizes a precision-recall curve as the mean of precisions at each threshold weighted by the change in recall since the last operating point:

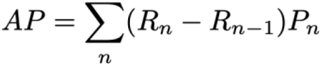

where *P_n_* and *R_n_* are the precision and recall at the n^th^ threshold. We report the model performance on the test data below:

| Class | Precision | Recall | F1-Score | Support |
| --- | --- | --- | --- | --- |
| 0 | 0.99 | 1.00 | 0.99 | 3159 |
| 1 | 0.97 | 0.89 | 0.93 | 251 |
| accuracy |  |  | 0.99 | 3410 |
| macro avg | 0.98 | 0.94 | 0.96 | 3410 |
| weighted avg | 0.99 | 0.99 | 0.99 | 3410 |

We also trained and evaluated five other classic machine learning classifiers including decision tree, gaussian naive bayes, logistic regression, support vector machine (SVM), and SVM with stochastic gradient descent for baseline methods comparison. While all of the methods tested achieved high accuracy score on the test set (>0.95), the random forest classifier outperforms other methods on both F1 and AP scores by a large margin.

All of the classifiers and evaluation metrics used in this study are implemented with the scikit-learn platform (Pedregosa et al., 2011).

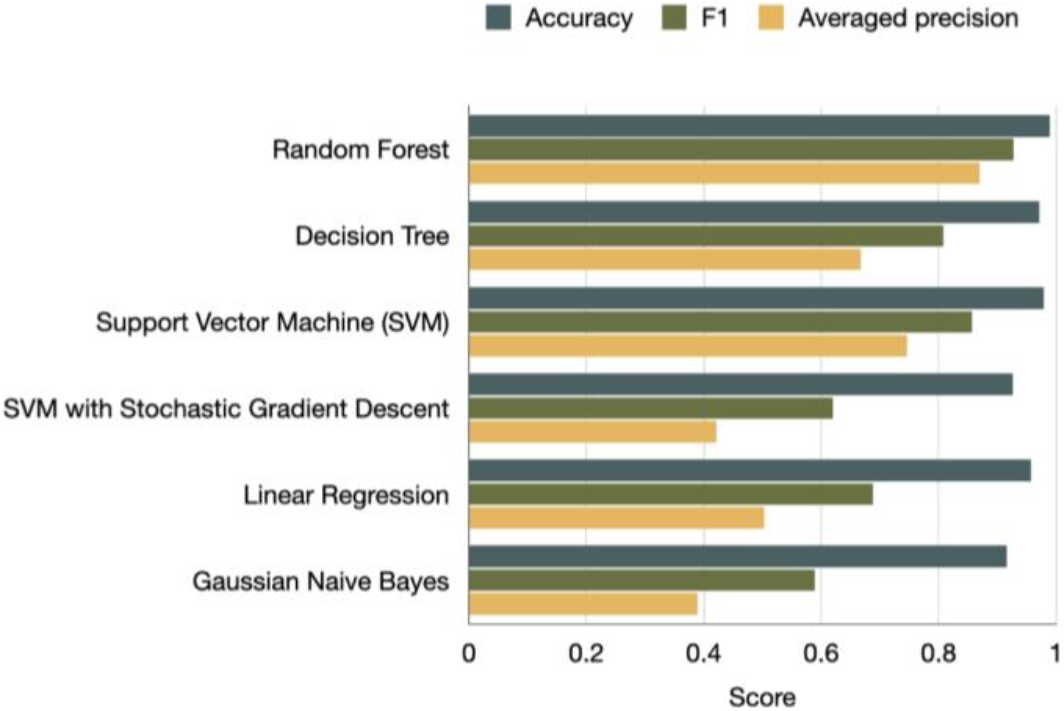

#### 1.5 TPC trainable option

In some cases, users may wish to re-train a TPC specifically tailored to their own dataset which can have distinctly different spindle dynamics and experimental design (e.g., imaging conditions, development stage, etc.). We propose that our framework is highly suitable for this purpose. A user may simply follow the steps detailed in our GitHub walk-through tutorial: https://github.com/yifnzhao/CentTracker, which includes feature construction, filtering, manual validation, hyperparameter tuning, model training, and evaluation. As demonstrated in Figure 3, our framework achieved strong performance in other *C. elegans* cell types and in cells from other species (here *Drosophila*).

